# Live imaging of Alu elements reveals non-uniform euchromatin dynamics coupled to transcription

**DOI:** 10.1101/2024.03.15.585287

**Authors:** Yi-Che Chang, Sofia A. Quinodoz, Clifford P. Brangwynne

**Author notes:** Correspondence: Clifford P. Brangwynne.

## Abstract

Chromatin structure and dynamics are crucial for eukaryotic nuclear functions. Hi-C and related genomic assays have revealed chromatin conformations, such as A/B compartments, in fixed cells, but the dynamic motion of such structures is not well understood. Moreover, elucidating the relationship between the motion of chromatin and transcriptional activity is hampered by a lack of tools for specifically measuring the mobility of active euchromatin. Here, we introduce a CRISPR-based strategy for live imaging of the gene-rich A compartment by labeling Alu elements — a retrotransposon family enriched within the transcriptionally active A compartment. Surprisingly, within euchromatin, microscopy analysis reveals that Alu-rich regions do not correlate with lower local H2B density, and form irregular foci of a few hundred nanometers in diameter, underscoring the heterogeneity of euchromatin organization. Alu-rich (gene-rich) chromatin is also more mobile than Alu-poor (gene-poor) chromatin, and transcription inhibition by actinomycin D results in decreased chromatin mobility of Alu-rich regions. These observations highlight the complexity of chromatin organization and dynamics and connect them to transcriptional activity on a genome-wide scale.

## Introduction

Genomic technologies have revealed complex chromatin organization at different length scales and linked this structure to potential functional roles. For example, at the megabase level, genomic contact mapping approaches (e.g., Hi-C) demonstrate that the human genome is divided into A and B compartments, each of which tends to interact with genomic loci within the same compartment (Lieberman-Aiden et al., 2009; Rowley et al., 2017). The A compartment is enriched in epigenetic marks associated with active transcription (Lieberman-Aiden et al., 2009), contains early-replicating regions (Marchal et al., 2019), occupies the interior of the nucleus (Kölbl et al., 2012), and largely overlaps with euchromatic areas devoid of densely packed heterochromatin (Rowley et al., 2017). Compartmentalization also changes during differentiation (Dixon et al., 2015; Zheng and Xie, 2019), accompanying gene expression and replication timing changes (Miura and Hiratani, 2022).

Despite these significant advances, genomic assays generally report on population-averaged structures and require chemical crosslinking. This limitation prohibits the measurement of chromatin dynamics in living cells, including spatiotemporal mobility of chromatin compartments, and how that relates to rapid processes occurring on the timescale of less than a minute, like transcription. Live-cell imaging methods have thus been crucial for studying real-time relationships between chromatin structure and its functions at the single-cell level. For example, CRISPR/Cas9-based genomic imaging and single-locus tracking have revealed a connection between chromatin mobility and the transcriptional activity of individual genes (Gu et al., 2018). However, it has been challenging to expand this understanding to a larger genomic context. While some studies have observed increased chromatin mobility associated with active transcription (Gu et al., 2018), contrasting behavior has also been reported (Ochiai et al., 2015). These single-locus tracking methodologies are very effective in measuring selected genes within specific biological contexts, but are less optimal for generating a broader understanding of specific chromatin structures, such as A/B compartments, due to their inherently low-throughput nature.

Approaches to labeling and tracking bulk chromatin allow for investigating chromatin dynamics on larger scales. For example, the application of particle image velocimetry (PIV) to time-lapse sequences of H2B images has revealed that chromatin displays micron-scale local cohesion over several seconds (Zidovska et al., 2013). Alternatively, rather than labeling nearly all chromatin, chromatin can be sparsely labeled for single-nucleosome tracking, well-suited for studying chromatin dynamics over shorter time and length scales (Xie and Liu, 2021). Nevertheless, these methods generally lack information regarding the local chromatin environment (e.g., epigenetic state) and genomic context (e.g., A/B compartments and TADs). Interpretations derived from the analysis of chromatin dynamics are thus challenging to integrate with chromatin structures mapped by Hi-C and other genomics studies, limiting our understanding of sequence-dependent chromatin mobility.

Here, we present a novel strategy for mapping the spatiotemporal dynamics of the A compartment of the human genome in living cells through sequence-specific chromatin labeling. This approach relies on targeting dCas9 to retrotransposon Alu elements, which are distributed throughout the genome but highly enriched in the A compartment, which is closely associated with transcriptionally active (Lieberman-Aiden et al., 2009) and more accessible regions of chromatin and is believed to coincide with euchromatin (Rowley et al., 2017). Our results reveal that while chromatin enriched in Alu elements is depleted in regions with high chromatin density, Alu density and chromatin density are not correlated in the less dense euchromatin. By integrating the live imaging of Alu elements with bulk chromatin labeling, we provide evidence that transcriptionally active chromatin spanning the A compartment exhibits increased mobility. Perturbations of transcription further revealed non-uniform impacts on chromatin dynamics. These results underscore the heterogeneity inherent within euchromatin.

## Results

### Targeting Alu-elements genome-wide enables live imaging of chromatin across the A compartment

We began by asking if there are repetitive sequences that can be targeted by dCas9 that would be representative of the majority of the A compartment. Bioinformatic analyses have shown that in the human genome, the A compartment is highly enriched in short interspersed nuclear elements, or SINEs (Lu et al., 2021), which constitute more than 10% of the human genome. The most abundant type of SINEs are Alu elements, a family of sequences averaging 300 base pairs in length and typically situated in gene-rich regions (Deininger, 2011; Lander et al., 2001). Indeed, sequences from Alu elements have previously been used as DNA FISH probes (Lu et al., 2021; Solovei et al., 2009) for imaging euchromatic regions of the genome, but this has been limited to fixed samples. We were thus encouraged by the potential to target and label the consensus sequence from Alu elements using a dCas9-based strategy. However, despite the repetitive and widespread nature of Alu elements, its linear density is not high. Thus, we decided to leverage existing signal amplification technologies, previously developed in CasDrop (Shin et al., 2019) and built upon SunTag (Tanenbaum et al., 2014), to develop an Alu-element live-imaging strategy (Figure 1 A; Figure S1 A). SunTag amplifies the fluorescence signal at target loci while keeping the dCas9-fusion construct short enough to be packaged by lentivirus.

**Figure 1:**
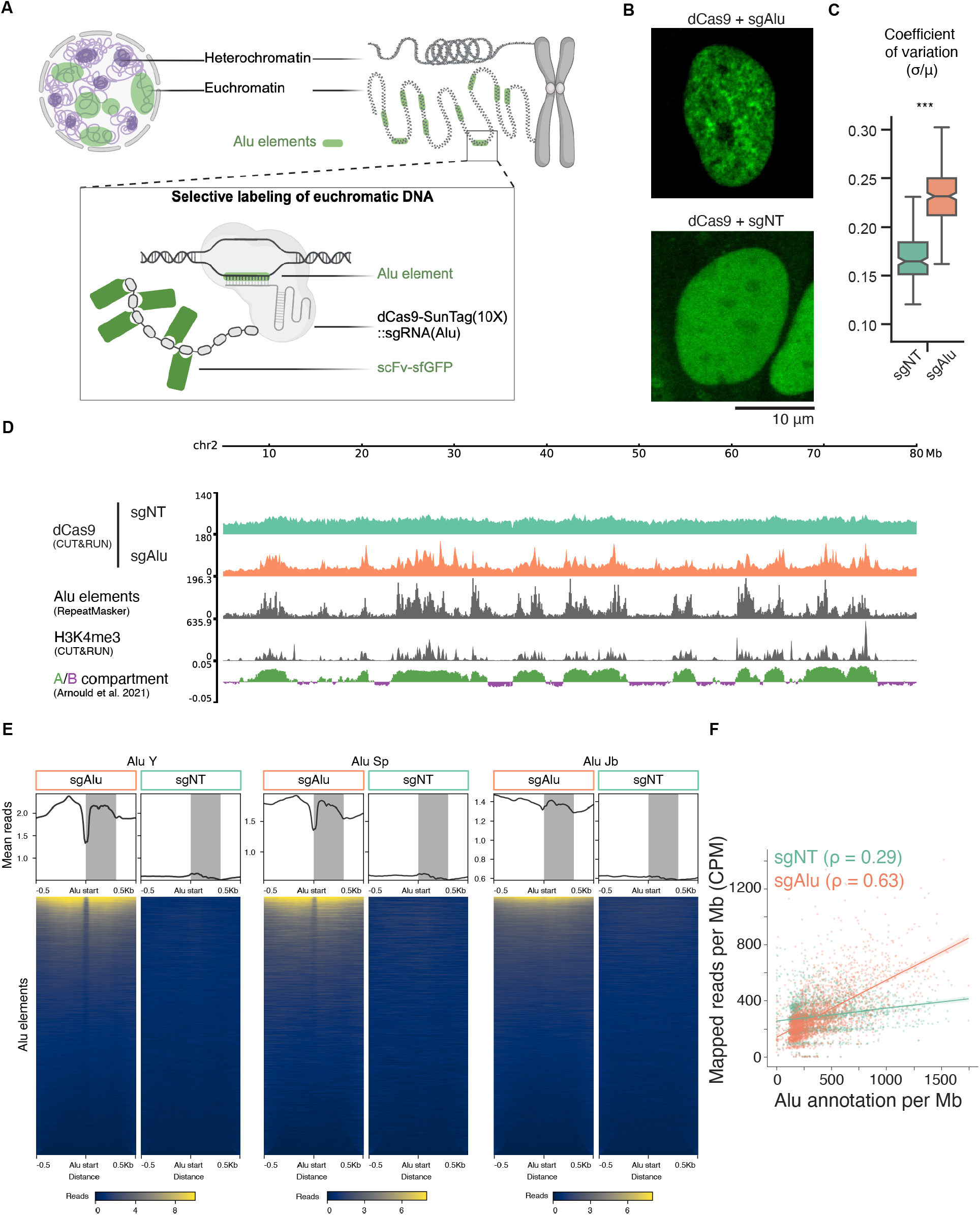
Live imaging of Alu elements with CRISPR/Cas9-based system. (**A**) Schematic illustrating how genome-wide Alu elements are targeted with dCas9 constructs. (**B**) Fluorescence images of U2OS cells labeled with Alu elements (sgAlu) or non-target control (sgNT) sgRNAs. Scale bar, 10 μm. (**C**) Coefficient of variance (standard deviation normalized by mean) of fluorescence intensity in nuclear pixels from (B). Notch represents s.d., box represents quartiles (lower, Q1; center, Q2; and higher, Q3), whiskers extend to data points that lie within 1.5 IQR (interquartile range = Q3 - Q1) of the lower (Q1) and higher (Q3) quartiles. *n* ≥ 300 nuclei for each group. *** denotes *P* < 0.001 using two-sided Brunner-Munzel test with t-distribution. (**D**) Genomic tracks of CUT&RUN sequencing assay against dCas9 compared to Alu-element annotations (from RepeatMasker), CUT&RUN against promoter-specific epigenetic mark (H3K4me3), and A/B compartments (A compartment > 0; B compartment < 0). For dCas9 CUT&RUN tracks, data ranges were scaled to account for total read counts for each condition. sgRNAs used in dCas9 CUT&RUN: control (sgNT) or Alu-targeting (sgAlu). A/B compartments were assigned from an existing Hi-C dataset from the same cell type (Arnould et al., 2021). Genomic range shown: chr2 5-80 Mb. (**E**) Averaged profiles and heatmaps showing mapped CUT&RUN reads at three Alu-repeat families annotated across the hg38 genome. (**F**) Relationship between density of sgNT or sgAlu-targeted dCas9 CUT&RUN reads across the genome per megabase and density of Alu annotations for the same window of 1 Mb. Spearman correlation coefficient: *ρ* = 0.63 for sgAlu, and *ρ* = 0.29 for sgNT.

**Figure S1:**
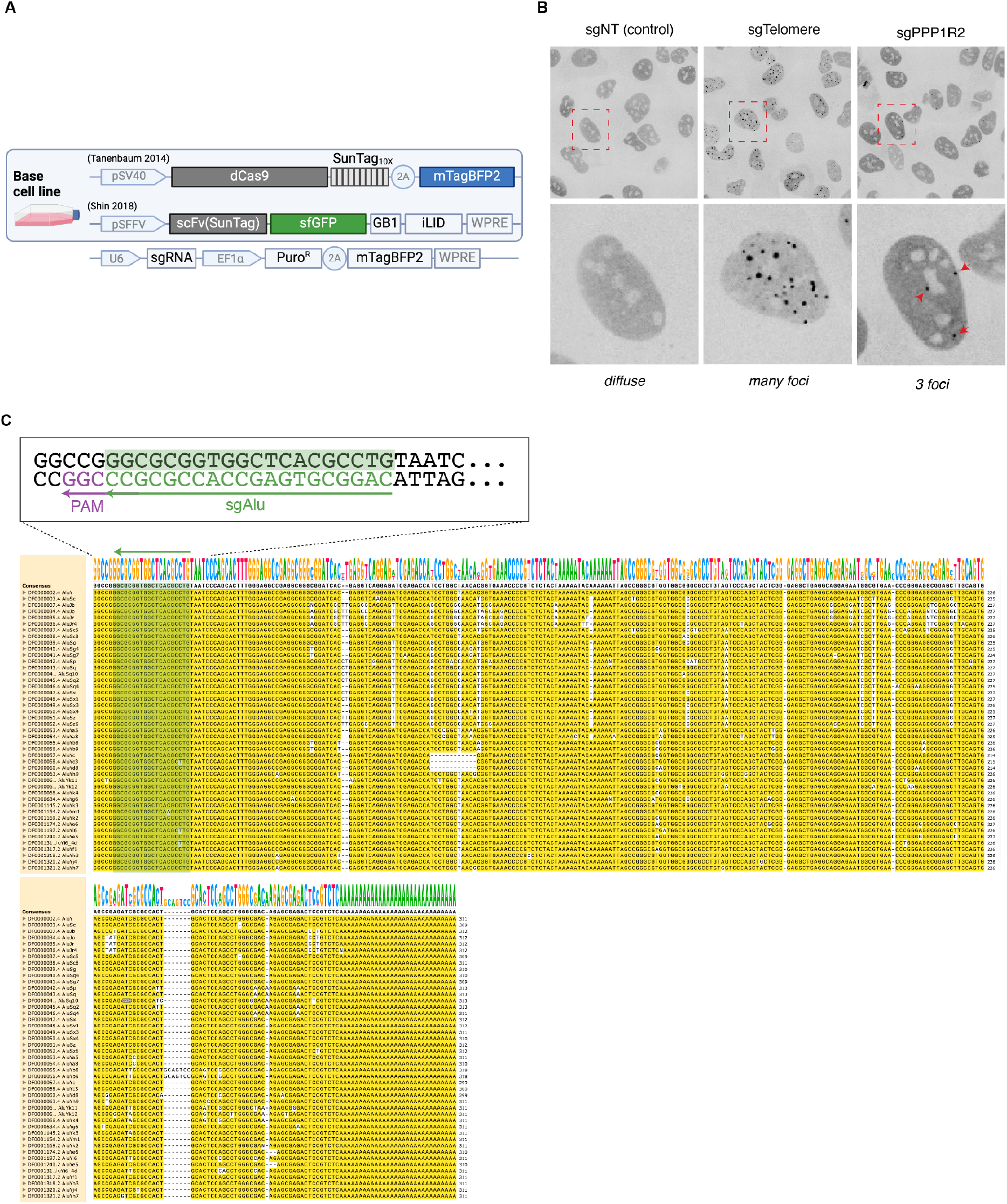
Optimized dCas9 cell line for targeting consensus sequence of Alu family. *Related to:* Figure 1. (**A**) Constructs for dCas9-based Alu-imaging. (**B**) Clonal line expressing dCas9-SunTag system optimized for genomic imaging. Fluorescence images from the dCas9 channel are shown. Repetitive targets include telomeres (sgTelomere) and *PPP1R2* (sgPPP1R2) as well as a non-target control (sgNT). (**C**) Alignment of Alu family sequences and sgAlu guide RNA target (green shaded). Zoomed-in: aligned sequence, and annotations for protospacer and PAM of sgAlu guide RNA design.

We optimized dCas9-expression level and generated clonal lines to image Alu elements. Their nuclear signal exhibits a textured pattern, distinct from the diffuse pattern observed in the non-target sgRNA control groups (Figure 1 B & C). We find that the expression levels of the specific components in the dCas9-based genomic imaging system are critical for imaging Alu elements with high signal-to-noise ratio. To this end, we generated clonal lines of U2OS cells expressing the dCas9-SunTag system stably at low levels. We identified desired clonal lines showing clear target foci for telomeres or the *PPP1R2* locus after transducing cognate sgRNA lentivirus for telomeric repeat sequence TTAGGG or an approximately 500-copy repeat that exists in the *PPP1R2* gene, respectively (Figure S1 B). These clonal lines were further screened by expressing a single sgRNA targeting 20 nucleotides of the 5’-end consensus sequence of Alu elements (Castanon et al., 2020) (Figure S1 C). A non-target sgRNA (sgNT) with non-human target (Castanon et al., 2020) was used as control throughout this study. The textured pattern of dCas9 observed is specific to Alu-targeting sequences (sgAlu) and is similar to previously described fixed-cell images of DNA FISH probing Alu elements (Bolzer et al., 2005; Lu et al., 2021).

To validate the specificity of our labeling approach, we mapped dCas9 binding using a CUT&RUN assay (Skene and Henikoff, 2017), which is akin to chromatin immunoprecipitation sequencing (ChIP-seq). Specifically, we performed CUT&RUN on a clonal cell line expressing the Alu-targeting or non-targeting sgRNA. CUT&RUN sequencing reads produced fragment size distributions comparable to those reported (Miura and Chen, 2020), with a major fraction between 120 to 270 bp (Figure S2 A). We find that dCas9 is highly enriched at Alu elements only when the cognate guide sgAlu is expressed and not when sgNT control is expressed (Figure 1 D). To measure whether dCas9 binding is enriched near actively transcribed loci, we performed CUT&RUN for H3K4me3, an epigenetic mark occurring at transcriptionally active promoters (Talbert et al., 2019), in the same cell line. We observe that Alu-targeted dCas9 localization correlates with H3K4me3 marks, suggesting dCas9 localization is enriched near sites of active transcription and within the A compartment (Figure 1 D), consistent with bioinformatic characterizations of Alu element distributions (Lander et al., 2001; Lu et al., 2021). To validate whether dCas9 binding is enriched at Alu elements genome-wide, we measured dCas9 binding across all annotated Alu repeat subfamilies. Specifically, we compared dCas9 enrichment at several Alu subfamily annotations and found that dCas9 binding is highly enriched at Alu-containing DNA sites in cells expressing sgAlu, but not in cells expressing sgNT control. As examples, the coverages around three subfamilies, Alu Y, Sp, and Jb, show enriched signal within annotated Alu element regions, and drop at the boundaries of those regions (Figure 1 E). The increased signal further away from Alu regions is likely due to adjacent Alu elements. Furthermore, there is a positive correlation between Alu-targeted dCas9 localization per megabase and the density of number of Alu annotations per megabase, confirming the specificity of Alu-element targeting genome-wide (*ρ* = 0.63) compared to non-targeted dCas9 localization (*ρ* = 0.29) (Figure 1 F). These clonal lines expressing the dCas9-SunTag system are thus suitable for live-cell imaging of the A compartment using a sequence-specific approach and are used throughout the rest of this study unless otherwise specified.

**Figure S2:**
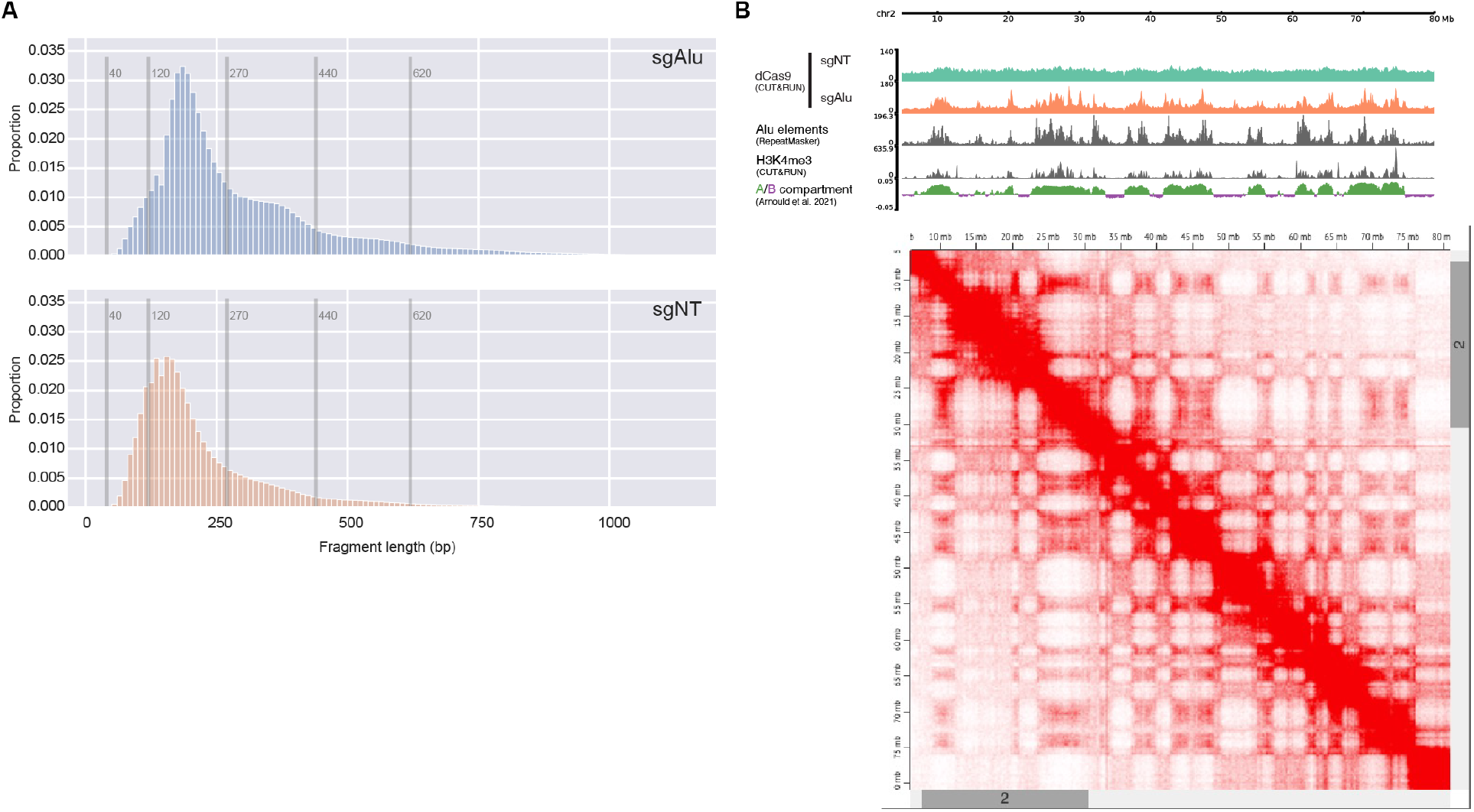
CUT&RUN sequencing verified the specificity of Alu targeting. *Related to:* Figure 1. (**A**) dCas9 CUT&RUN library insert size post adapter trimming for (top) sgAlu and (bottom) sgNT. (**B**) U2OS Hi-C contact matrix on chromosome 2 (5-80Mb) alongside genomic tracks shown in Figure 1 D. Hi-C data obtained from an existing dataset (Arnould et al., 2021).

### Alu elements do not correlate with histone density within euchromatin

Our dCas9-SunTag approach to label Alu-rich regions of the genome enables us to probe the localization of chromatin across the A compartment in live cells. We began by comparing the Alu pattern to other subnuclear structures that Alu elements are known to be enriched or depleted at, such as nuclear speckles and heterochromatin, respectively. Specifically, we expressed fluorescent protein miRFP670-fusion of markers of these structures in the clonal line, and transduced lentivirus for either sgRNA targeting the consensus Alu element sequence (sgAlu) or non-target control (sgNT) sgRNA. We then imaged these dual-labeled cells, and analyzed their spatial patterns (Figure 2 A, B; see Methods). We found that Alu elements are generally depleted in dense constitutive heterochromatin regions labeled by HP1*α*, consistent with Alu depletion in heterochromatin (Lu et al., 2021). By contrast, Alu elements and nuclear speckles, labeled by SRRM1, have a higher correlation, consistent with previous findings showing Alu elements are enriched around nuclear speckles and genes of high transcriptional activity (Chen et al., 2018; Su et al., 2020).

**Figure 2:**
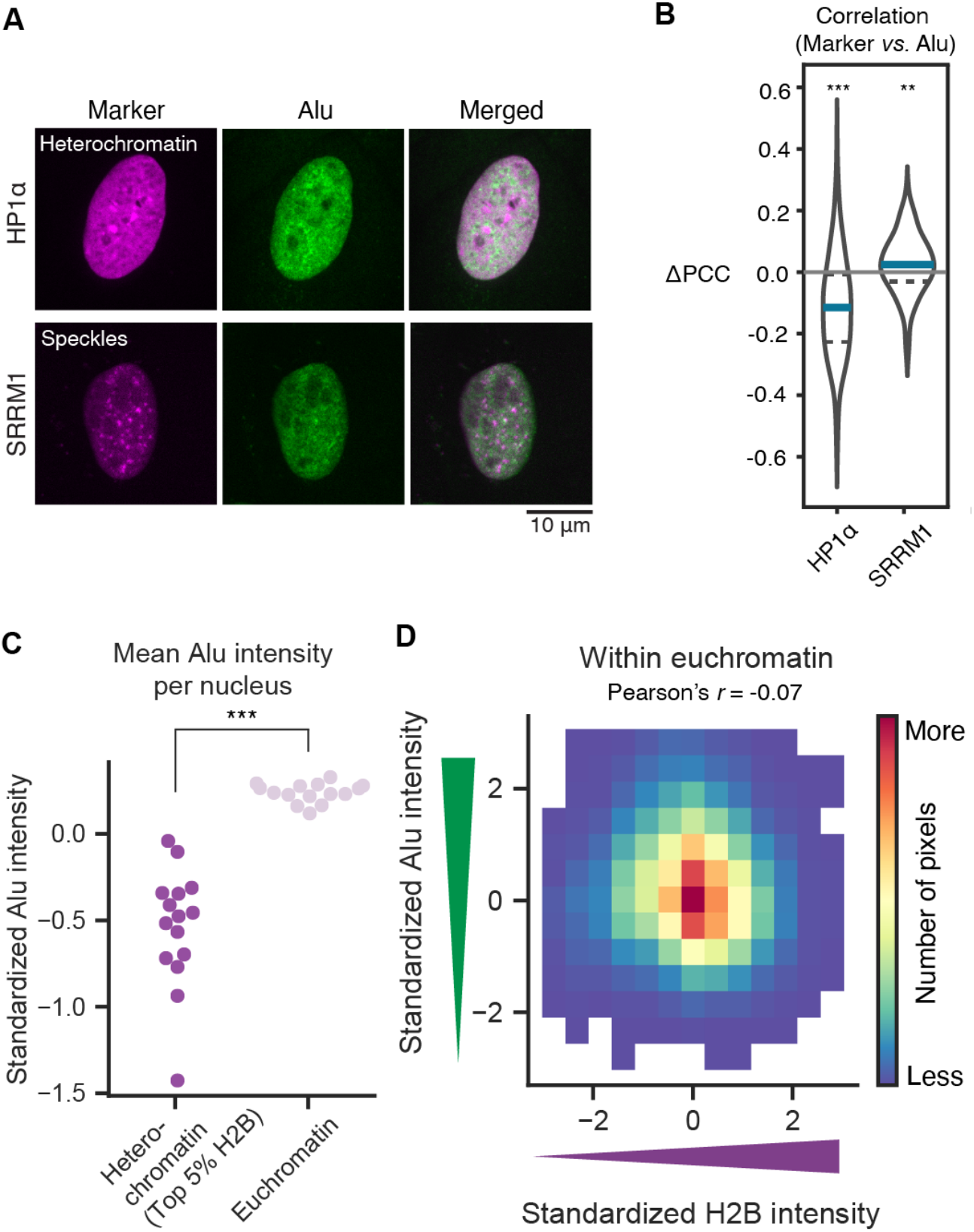
Alu-rich regions are depleted in heterochromatin but Alu elements are not correlated with histone density within euchromatin. (**A**) Fluorescence images of Alu-labeled cells co-expressing markers of distinct subnuclear regions: HP1*α* for heterochromatin and SRRM1 for nuclear speckles. Images are shown in individual and merged channels (magenta: marker; green: Alu elements) as annotated. Scale bar, 10 μm. (**B**) Change in Pearson correlation coefficient (PCC) of pixel intensities in nucleoplasmic regions between the marker and Alu elements (Alu-targeted dCas9) channels in (A), compared to the mean of control sgRNA (sgNT). Data is shown as violin plots (estimated probability density) with median (blue solid line, middle) and first and third quartiles (dashed lines, bottom, and top, respectively) inside. Grey solid line: change = 0 (mean of PCC for sgNT). *n* ≥ 150 nuclei for each group. ** denotes *P* < 0.01 and *** *P* < 0.001 using two-sided Brunner-Munzel test with t-distribution to compare sgAlu and sgNT conditions for each marker. (**C**) Mean standardized Alu intensity in euchromatin and heterochromatin regions. Each dot corresponds to a nucleus. *n* = 15 nuclei. *** denotes *P* < 0.001 using Mann-Whitney U rank test. See Figure S3 A for Alu- and H2B-intensity distributions in an example cell. (**D**) Joint distribution of Alu and H2B pixel intensities within euchromatin. Pearson correlation coefficient *r* = −0.07.

We next considered the relationship between Alu elements and histone density. First, we expressed H2B-emiRFP670 in the Alu-imaging clonal line, and compared the distribution of Alu intensity in regions with low or high histone density. The histone-dense regions, or heterochromatin, were segmented based on H2B intensity (top 5% H2B pixel intensity; see Methods), and we defined the euchromatin regions in our images by excluding heterochromatin and nucleolar areas. To enable comparison across various nuclei, pixel intensities were normalized (see Methods). We find that the mean Alu intensity per cell is significantly higher in euchromatin than in heterochromatin (Figure 2 C & Figure S3 A). This is consistent with previous observations that Alu elements tend to be biased towards the A compartment, which is typically thought to overlap with euchromatin, genome-wide (Chen et al., 2018; Lu et al., 2021). However, within the euchromatin region, we find a negligible correlation between Alu intensity and H2B intensity (*r* = −0.07) (Figure 2 D & Figure S3 B). This suggests that Alu-element density and histone density each provide potentially complementary information – Alu elements as chromatin identity (gene-rich vs -poor) and histone density as chromatin environment (compact or open) – that can be utilized to differentiate local chromatin contexts when analyzing chromatin behavior with subnuclear resolution.

### Alu-element density positively correlates with euchromatin mobility

We next asked if these gene-dense and gene-poor regions, identified by Alu-rich and Alu-poor regions, respectively, have unique chromatin dynamics. Because we can simultaneously image Alu elements and H2B in the same cell, we leveraged our Alu-element labeling to spatially encode genomic identity (Alu-rich or Alu-poor) as a context for H2B chromatin dynamics (Figure 3 A, B, C). This allows us to test if there is chromatin context-dependent H2B mobility behavior within the euchromatin region (Figure 3 D & Figure S4 A, B). For example, we can ask if H2B (chromatin) displacement is larger in the Alu-rich area than in the Alu-poor area.

**Figure 3:**
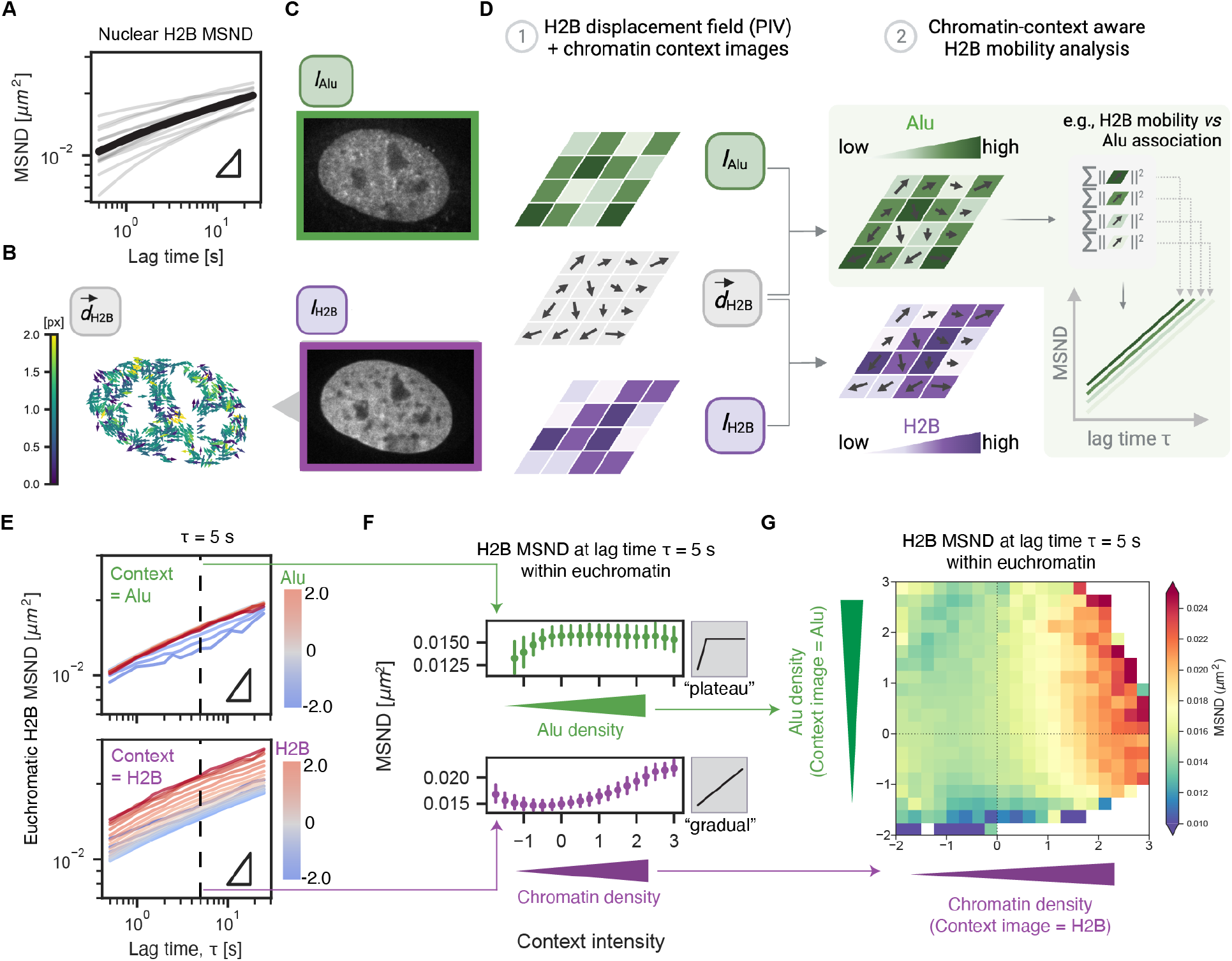
Context-aware histone mobility analysis integrating local Alu-element intensity uncovers sequence-specific mobility across euchromatin. (**A**) Chromatin MSND in euchromatin regions plotted against lag time. Data is shown as mean (thick line) over *n* = 15 individual nuclei (thin lines). Triangle represents slope = 0.5. (**B**) An H2B displacement field corresponds to the nucleus shown in (C). Displacement vectors from nucleoli and heterochromatin regions are excluded (see Methods). (**C**) Representative images for Alu and H2B channels. (**D**) Framework for context-aware chromatin mobility analysis. H2B displacement fields are spatially combined with context image(s), allowing for the analysis of chromatin mobility in different chromatin context(s). (**E**) H2B MSND plotted against lag time and stratified against respective chromatin context (Z-score): Alu density (*top*) and H2B density (*bottom*). Colors represent relative context image intensity. Triangles represent slope = 0.5. (**F**) Dependence of H2B MSND at lag time τ = 5 s on chromatin contexts (Z-score): Alu density (*top*) and H2B density (*bottom*). (**G**) Heat map showing H2B MSND at different combinations of Alu density (Z-score) and H2B density (Z-score), at lag time τ = 5 s. Colors represent the squared displacement in μm^2^. In (E), (F), and (G), *n* = 15 nuclei.

**Figure S3:**
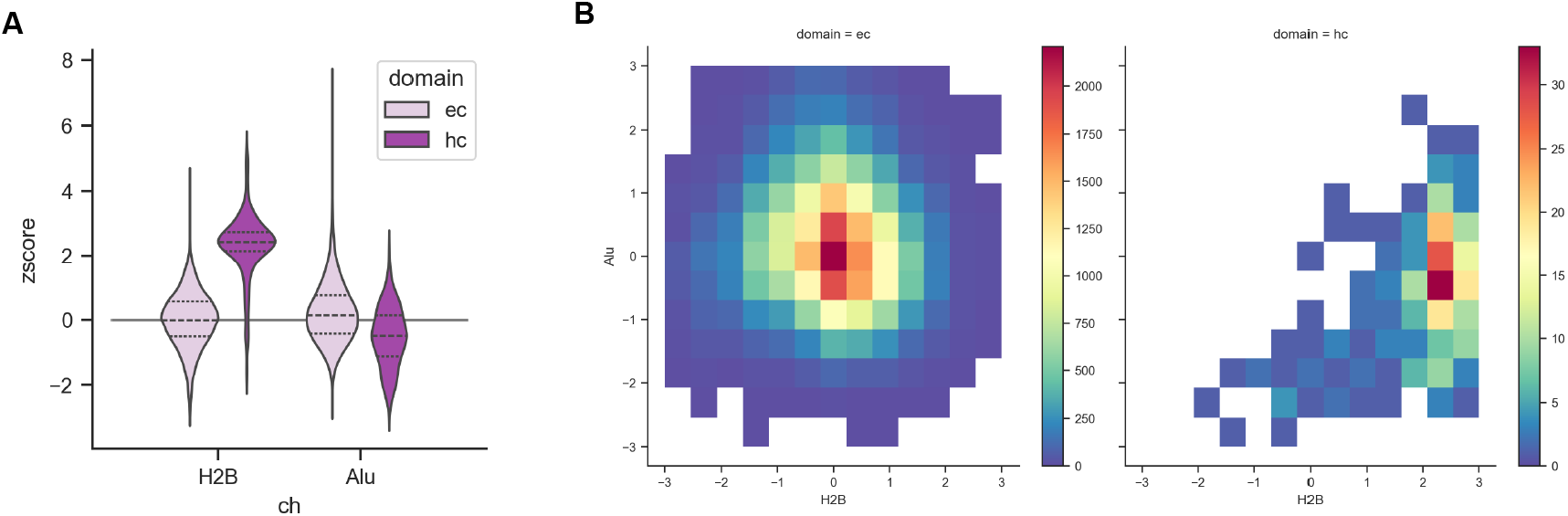
Relationship between Alu and H2B pixel intensities in euchromatin and heterochromatin regions. *Related to:* Figure 2. (**A**) Mean standardized H2B (*left*) and Alu (*right*) intensity distributions in euchromatin (*light*) and heterochromatin (*dark*) regions in a single cell provided as an example. (**B**) Joint distribution of Alu and H2B pixel intensities within either euchromatin or heterochromatin regions. Pearson correlation coefficients: *r* = −0.07 for euchromatin, and *r* = 0.35 for heterochromatin. The Euchromatin panel is the same as Figure 1 and is shown here for direct comparison with the heterochromatin panel.

**Figure S4:**
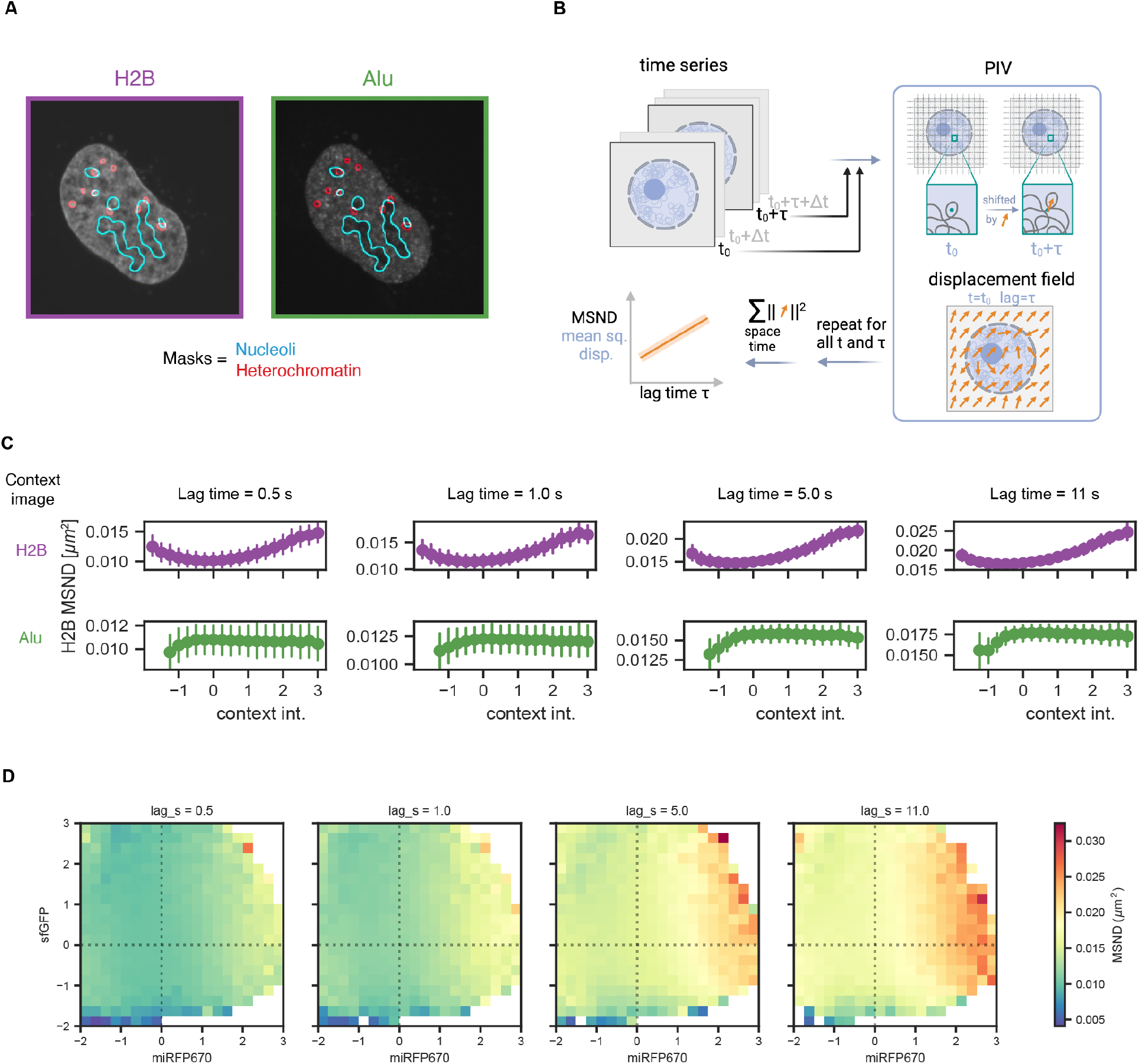
Details for PIV workflow and H2B MSND at different lag times. *Related to:* Figure 3. (**A**) Representative images of H2B and Alu channels and the masks for nucleoli and heterochromatin regions used in MSND analyses. (**B**) Schematics illustrating PIV workflow to estimate chromatin network displacement field(s). (**C**) Dependence of H2B MSND at different lag time τ on chromatin contexts: H2B density (*top*) and Alu density (*bottom*). Lag times τ = 0.5 s, 1.0 s, 5.0 s and 11.0 s are shown. (**D**) Heat maps showing H2B MSND at different combinations of Alu density (sfGFP) and H2B density (miRFP670) at different lag times τ = 0.5 s, 1.0 s, 5.0 s and 11.0 s. Colors represent the squared displacement in μm^2^. In (C) and (D), *n* = 15 nuclei.

To measure chromatin dynamics, interphase chromatin was labeled by expressing histone protein H2B tagged with fluorescent protein miRFP670 in the optimized Alu-imaging cell line (Figure 3 C). Movies of nuclei were recorded every 0.5 s for 60 s, using two cameras for capturing images in both channels simultaneously. We then measured chromatin network dynamics using particle image velocimetry (PIV), previously used to report chromatin dynamics from fluorescent histone H2B protein images (Zidovska et al., 2013). For each pair of two images from one such movie, PIV was applied to extract the displacement field; the resulting displacement field corresponds to a specific lag time τ = *Δt*, which is the difference in time between the two chosen frames (Figure S4 B). We excluded displacement vectors from nucleoli, nuclear envelope, and heterochromatin regions (Figure S4 A) in our analysis because PIV is sensitive to sharp changes in intensity around these boundaries. Repeating these procedures for all possible pairs and all accessible lag times yields displacement fields 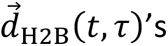 whose average — mean square network displacement (MSND) — reports the network dynamics (Figure S4 B). Our measurements of total chromatin dynamics at lag time τ = 5 s are typically on the order of 10^-2^ μm^2^ (Figure 3 A, B), in agreement with past studies (Shaban et al., 2020; Zidovska et al., 2013).

To compare if gene-rich and gene-poor (Alu-rich and Alu-poor, respectively) chromatin have different mobility, we focus on the MSND at lag time τ = 5 s to compare the measured displacement. This is the timescale over which mesoscale chromatin structure (across several μm) has been observed to have coherent movement (Zidovska et al., 2013). We observed an approximate power law relation between resulting H2B MSND (at regions with different Alu-element density or H2B density) and lag time (Figure 3 E), validating sufficient statistics obtained with our context-aware analysis framework. As Alu intensity increases, chromatin MSND increases and then plateaus, suggesting the average chromatin mobility is similar above a certain level of Alu density (Figure 3 F). This suggests that Alu-rich chromatin has a higher mobility than Alu-poor chromatin. Indeed, direct tracking of individual Alu foci with enhanced spatial resolution (Figure S5 A) suggests an anomalous exponent close to 0.5 (Figure S5 B), consistent with subdiffusive behavior reported for gene loci (Gu et al., 2018). Additionally, applying PIV and MSND analysis to Alu signal also showed similar behavior within the same timescale (0.5 s ≤ τ ≤ 1 s) (Figure S5 C). Conversely, we observed a more gradual transition in chromatin MSND with respect to chromatin density. Interestingly, high histone density within euchromatin regions (defined as whole nucleus excluding heterochromatin and nucleolus, as described earlier) is associated with higher mobility (Figure 3 F & Figure S4 C). This is consistent with recent studies showing that euchromatin can be condensed while maintaining its high mobility (Maeshima et al., 2023; Miron et al., 2020; Nozaki et al., 2023). To further investigate the dependencies and interactions between H2B density and Alu intensity, we created a two-dimensional map of chromatin mobility (MSND) (Figure 3 G & Figure S4 D). The horizontal and vertical axes represent H2B density and Alu intensity, respectively. This analysis confirmed the trends from the one-dimensional context-dependent mobility analysis (Figure 3 F). Taken together, our data suggests that Alu-rich (gene-rich) regions have increased chromatin mobility compared to Alu-poor (gene-poor) regions.

**Figure S5:**
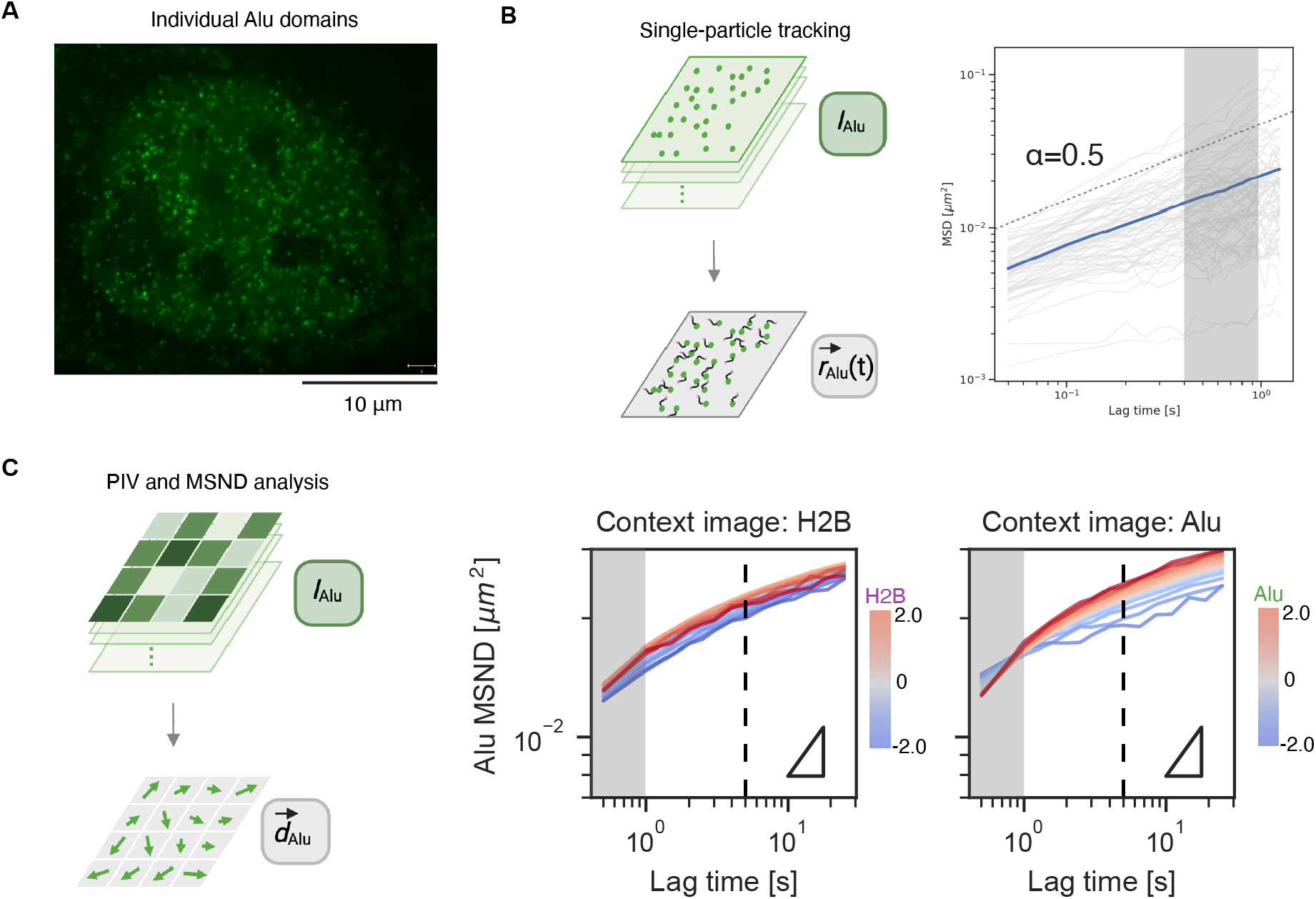
Alu-element mobility probed by single-particle tracking and PIV. *Related to:* Figure 3. (**A**) Fluorescence image of higher resolution, compared to Figure 1 B, allowing visualization of individual Alu-element foci. Scale bar, 10 μm. (**B**) MSD plotted against lag time for single particle tracking of Alu-elements domains. Data shown as time-averaged MSD (grey) and their mean (blue). (**C**) Alu MSND plotted against lag time and stratified against respective chromatin context: Alu density (*right*) and H2B density (*left*). Colors represent relative context image intensity (Z-score). Triangles represent slope = 0.5. *n* = 15 nuclei. Figure 3 E is the H2B MSND equivalent. Shaded areas in (B) and (C) correspond to the same range of lag time.

### Changes in chromatin mobility upon transcription inhibition are context-specific

Given that A-compartment regions are enriched in Pol II occupancy and histone acetylation marks on a genome-wide scale (Saxton et al., 2023) and that these gene-dense (Alu-rich) regions exhibit higher mobility than gene-poor (Alu-poor) regions (Figure 3), we hypothesized that the dynamics of Alu rich chromatin are coupled to Pol II transcriptional activity. To test this, we investigated how these unique regions of the genome respond to perturbations of Pol II transcription.

We treated cells for 4 to 6 hours with a panel of Pol II transcription inhibitors previously reported (Ku et al., 2022; Nagashima et al., 2019; Zidovska et al., 2013) to affect nucleosome mobility: *α*-amanitin (aAM), flavopiridol (FVP), and actinomycin D (ActD) (Figure 4 A). We first examined the mean chromatin displacement across the entire euchromatin compartment. Chromatin mobility is either not affected or may become slightly more mobile (by 10-20%) upon inhibition of Pol II transcription using *α*-amanitin and flavopiridol (Figure 4 B; Figure S6 A), comparable to results from both bulk-chromatin and single-nucleosome tracking (Nagashima et al., 2019; Zidovska et al., 2013). In contrast, actinomycin D, which blocks both Pol I and II transcription at the applied concentration of, significantly slows down chromatin dynamics by approximately 40% (Figure 4 B; Figure S6 A), consistent with previous studies (Nagashima et al., 2019; Zidovska et al., 2013).

**Figure 4:**
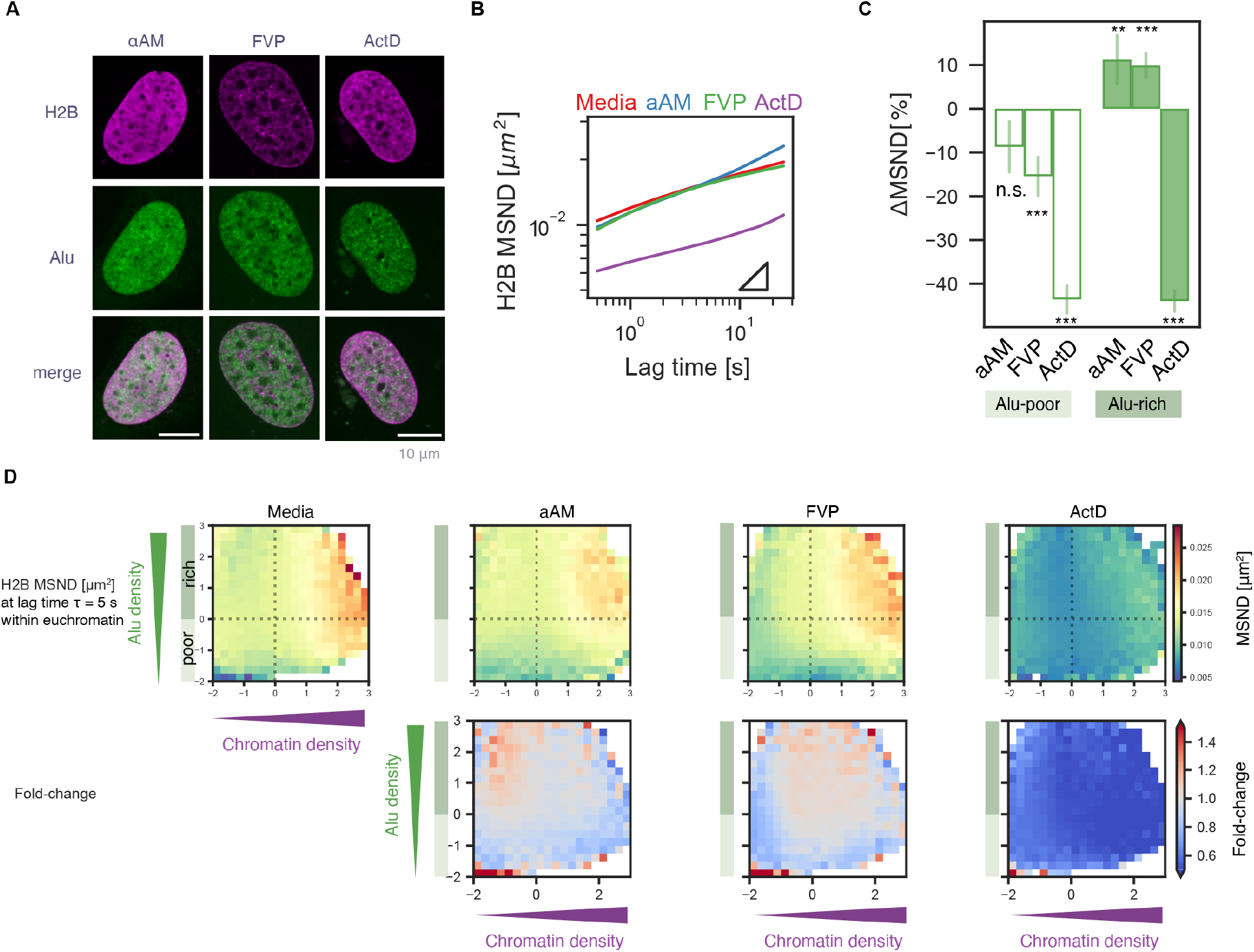
Alu element-specific chromatin mobility is reduced upon Pol II transcription inhibition. (**A**) Representative fluorescence images of U2OS cells treated with transcription inhibitors *α*-amanitin (aAM), flavopiridol (FVP), and actinomycin D (ActD) for individual channels or merged. *Magenta*, H2B; *green*, Alu elements. Scale bars, 10 μm. (**B**) H2B MSND euchromatin region plotted against lag time for each treatment. Triangles represent slope = 0.5. (**C**) Percent change in H2B MSND, at lag time τ = 5 s, in Alu-rich or Alu-poor regions after treatment. Data represented as mean ± s.e.m. ** denotes *P* < 0.01, *** *P* < 0.001, and n.s. not significant using two-sided Brunner-Munzel test with t-distribution to compare each treatment to the control (media). (**D**) Heat maps showing (*top*) H2B MSND at different combinations of Alu density and H2B density, at lag time τ = 5.0 s, after treatment, and (*bottom*) corresponding fold-change compared to control (media). Colors represent (*top*) squared displacement, in μm^!^, and (*bottom*) fold change. For (B), (C), and (D), *n* ≥ 10 nuclei for each condition.

**Figure S6:**
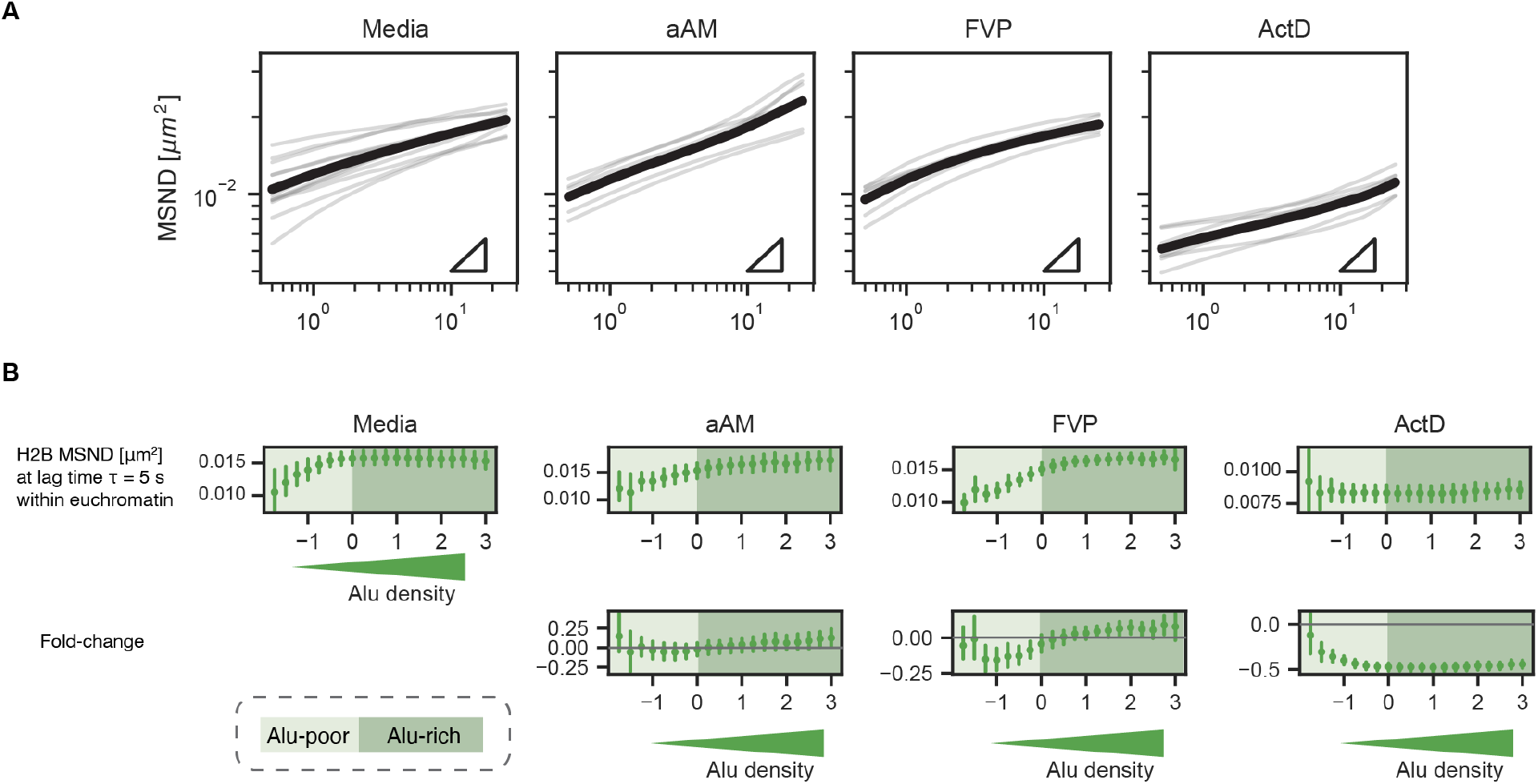
H2B MSND behaviors upon Pol II transcription inhibition. *Related to:* Figure 4. (**A**) Individual ensemble MSND traces for each nucleus are shown (light) together with the corresponding mean (heavy). Triangles represent slope = 0.5. Related to Figure 4 B. (**B**) (*Top*) H2B MSND at different degrees of Alu density (Z-score), at lag time τ = 5 s, after each treatment, and (*bottom*) corresponding fold-change (compared to media-only). Alu-rich (dark-shaded) or -poor (light-shaded) regions are separated at standardized Alu intensity (Z-score) = 0. Data represented as mean ± s.d. For (A) and (B), *n* ≥ 10 nuclei for each condition.

We next used the context-aware approach to analyze the mobility of sub-categories of chromatin upon transcription inhibition. When we compared *α*-amanitin- and flavopiridol-treated conditions to the control (media), H2B MSND at lag time τ = 5 s in the Alu-rich regions showed a modest increase in MSND, by approximately 10% (Figure 4 C, D; Figure S6 B). Conversely, transcription inhibition induced by actinomycin D slows down the entire analyzed euchromatin region by approximately 40%, irrespective of its Alu density (Figure 4 C, D; Figure S6 B). These results highlight the complexity of chromatin behavior and emphasize the importance of considering multiple factors that describe the identity and environment of chromatin locally.

## Discussion

Chromatin dynamics are intimately coupled to genome organization and function. Our study sought to characterize the dynamic mobility and organization of active chromatin in living cells by microscopy. Previously, chromatin organization has primarily been examined in fixed cells using crosslinking-based genomic techniques like Hi-C (Lieberman-Aiden et al., 2009) or in a limited capacity in live cells through individual locus labeling (Chen et al., 2013; Gu et al., 2018; Ku et al., 2022; Ochiai et al., 2015). We introduce a flexible strategy that targets dCas9 to Alu elements, which are ubiquitous and widely distributed within euchromatic chromatin regions, to characterize the dynamics of euchromatic regions in living cells. In contrast to live tracking techniques that focus on selected gene loci, our approach allows imaging of an entire class of chromatin structure while maintaining sequence-specificity.

Our simultaneous imaging of Alu elements and histone protein H2B revealed that the density of Alu elements is decoupled from histone density in non-heterochromatic regions. This is surprising, as transcriptionally active euchromatic regions are generally characterized to be anti-correlated with chromatin density, but agrees with recent studies where densely packed nucleosome domains were observed in regions with transcriptionally active marks (Miron et al., 2020; Nozaki et al., 2023). This decoupled view of H2B density (as a proxy of chromatin density) and Alu-element density (as a proxy of gene density) is also consistent with recent studies identifying high density chromatin regions within “euchromatin” regions. ATAC-PALM demonstrated that transposase-accessible chromatin forms spatially segregated domains that are 150 nm in diameter and transcriptionally active (Xie et al., 2020). Labeling individual nucleosomes in early-replication foci combined with single nucleosome tracking suggested that histones in euchromatin form condensed solid-like structures of similar size (Nozaki et al., 2023). Our Alu-element imaging also reveals distinct domains of comparable size, and they exhibit subdiffusive behavior with an anomalous exponent of around 0.5 in living cells (Figure S5 B). These lines of evidence suggest that while euchromatin can be, on average, more open than heterochromatin, it is nevertheless heterogeneous in its internal organization.

We observed that chromatin enriched in Alu elements are either unaffected or may become slightly more mobile (Figure 4) when transcription is perturbed by *α*-amanitin, which degrades Pol II (Bensaude, 2011), and flavopiridol, which inhibits CDK9 (Bensaude, 2011), implying that machinery involved in active transcription could slow down chromatin dynamics. The change in mobility of Alu-rich chromatin upon Pol II transcription inhibition is unlikely to be caused by the transcription of Alu elements themselves, which are transcribed by Pol III and only at a low level (Liu et al., 1994; Paulson and Schmid, 1986). Instead, chromatin mobility in Alu-rich areas most likely reflects the mobility of Pol II-regulated genes, a lot of which contains intronic Alu elements. Increased chromatin mobility in Alu-rich regions upon flavopiridol and and *α*-amanitin has been reported inprevious studies, which suggested that Pol II transcription can slow down chromatin mobility (Nagashima et al., 2019; Ochiai et al., 2015; Zidovska et al., 2013). Interestingly, transcription inhibition by actinomycin D resulted in the opposite effect: a decrease in chromatin mobility globally across the nucleus, irrespective of the degree of Alu density (Figure 4). This effect may arise from accumulation of stalled polymerases (Kimura et al., 2002), which could slow down overall chromatin dynamics. A recent simulation-based study has proposed a unified view on the opposite effects of transcription inhibition on chromatin dynamics, wherein polymerases exert forces on chromatin and thus slow down its dynamics (Shin et al., 2023).

While polymerase activity clearly impacts chromatin dynamics, various processes directly or indirectly associated with transcription could also influence chromatin dynamics and contribute to the overall higher chromatin mobility in the Alu-rich areas. These include chromatin remodeling and loop extrusion, epigenetic modifications, changes in compaction states, and tethering to subnuclear structures like the nuclear lamina and nuclear speckles. Although we have successfully investigated chromatin dynamics, focusing on the interplay between Alu elements and histone density, there remains a possibility that we were blind to some of these factors. These non-monitored factors might explain the different impacts of transcriptional inhibition on chromatin mobility, observed here (Figure 4) and previously reported in the literature.

In conclusion, our results on chromatin dynamics reveal that (1) H2B density alone, which is typically used as a measure of chromatin density, does not adequately differentiate mobility in Alu-rich and -poor areas with similar chromatin density, and (2) the impact of transcription on chromatin mobility is not uniform across different chromosomal contexts: Alu-rich chromatin may be more sensitive upon flavopiridol and *α*-amanitin treatments compared to Alu-poor chromatin (Figure 5). It could prove insightful to simultaneously monitor a set of chromatin contexts (e.g., epigenetic marks or specific DNA sequences) and the mobility of chromatin itself. Future work combining CRISPR-based labeling, histone PTM imaging in living cells (Sato et al., 2013; Saxton et al., 2023), and chromatin dynamics measurement would help comprehensively characterize the relationship between chromatin mobility and transcriptional activity. It is also possible to leverage iLID (Guntas et al., 2015; Shin et al., 2019), the optogenetic module already part of our labeling implementation, to bring in epigenetic writers to induce targeted chromatin remodeling (Bintu et al., 2016; Eeftens et al., 2021) and measure their effect on chromatin dynamics.

**Figure 5:**
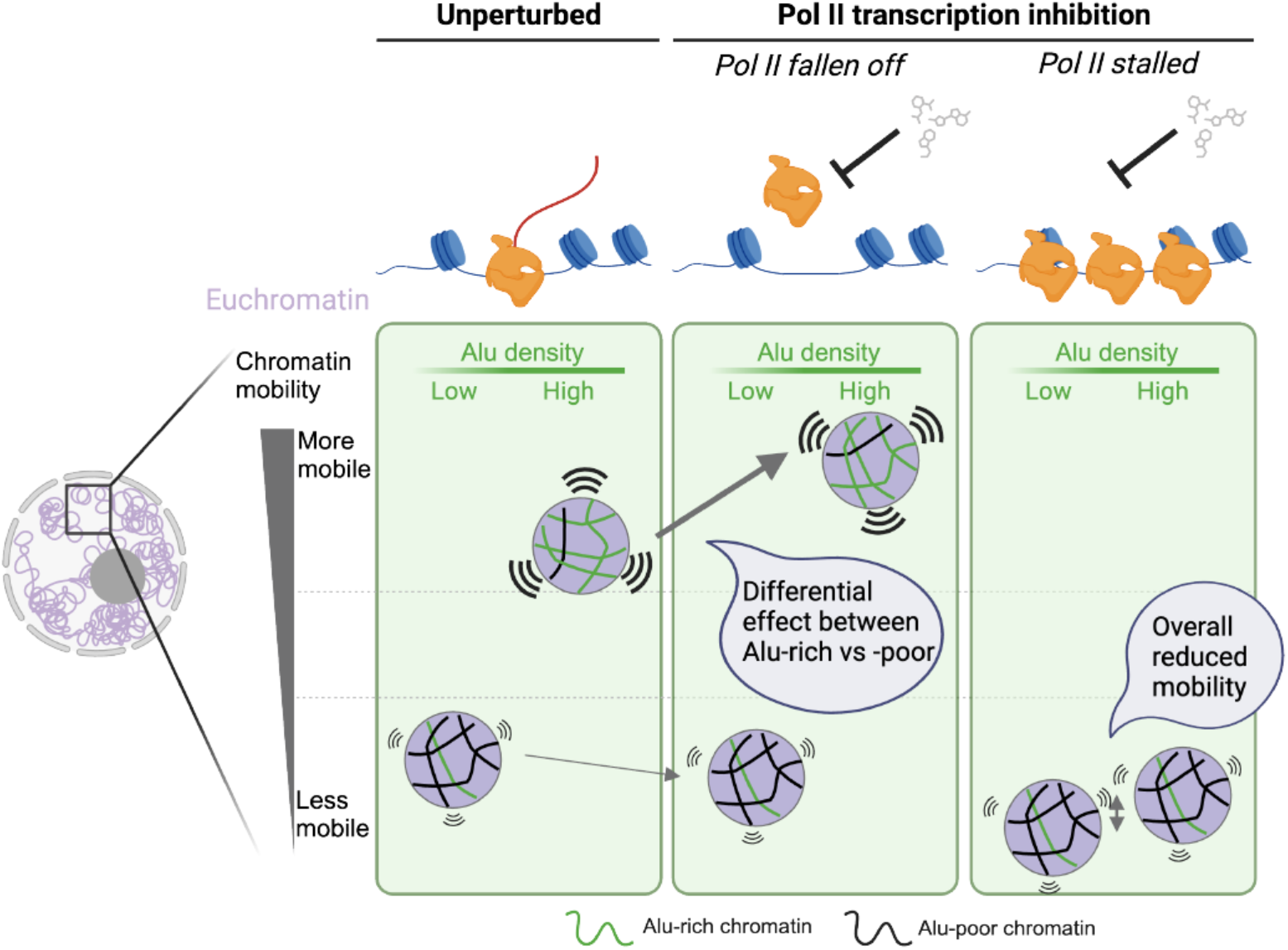
Distinct euchromatic sub-domains with higher mobility coupled to transcription. Schematic illustrating the non-uniform changes in euchromatin mobility between Alu-rich and -poor areas. Mobility of Alu-rich chromatin might increase upon Pol II transcription inhibition, where transcription machinery falls off from chromatin (e.g., *α*-amanitin and flavopiridol), and might decrease if transcription machinery gets stalled on chromatin (e.g., actinomycin D).

### Limitation of study

Our live-cell A-compartment imaging is powerful as it targets and labels a ubiquitous sequence genome-wide. Nevertheless, we cannot rule out the possibility of changes in transcriptional activity upon recruiting dCas9 onto chromatin at potentially thousands of Alu elements. That said, other work tracking gene loci, such as promoter or enhancer regions, using dCas9 demonstrated no significant change in the transcriptional activity of genes (Gu et al., 2018). Because Alu-element distribution is correlated with promoters and enhancers (Lu et al., 2021, 2020), we assume dCas9 targeting in this study similarly has minimal impact on transcription and chromatin mobility.

## Acknowledgments

We thank Aaron E. Lin and Britt Adamson for insightful discussions on CRISPR/Cas9-based technology optimization; Amy R. Strom, Ushnish Rana, and Mackenzie T. Walls for critical reading and comments on the manuscript; Daniel S.W. Lee for early discussions; Lennard W. Wiesner and Jessica Z. Zhao for materials; and Evangelos G. Gatzogiannis for assistance with imaging. We thank all current and previous members of Brangwynne Lab for helpful discussions and feedback.

We thank C. DeCoste, K. Rittenbach, G. Palmieri, and the Molecular Biology Flow Cytometry Resource Facility, partially supported by the Rutgers Cancer Institute of New Jersey NCI-CCSG P30CA072720-5921. We thank the Princeton University Genomics Core Facility Staff for Illumina sequencing. Some illustrations were created with BioRender.com.

S.A.Q. is funded by the HHMI Hanna H. Gray Fellowship. This work was funded by the Princeton Center for Complex Materials, an NSF MRSEC (DMR-2011750); the AFOSR MURI (FA9550-20-1-0241); the St. Jude Research Collaborative on the Biology and Biophysics of RNP granules; and the Howard Hughes Medical Institute.

## Author Contributions

Y.-C. C., S.A.Q., and C.P.B. conceptualized the study. Y.-C. C. and S.A.Q. performed experiments and formal analyses. Y.-C. C., S.A.Q., and C.P.B. wrote the paper. C.P.B. acquired the funding for this work.

## Materials and Methods

### Sample preparation

#### Plasmids

All plasmids used in this study are listed under the *Plasmids* section in Table 1. sgRNAs for *PPP1R2* (sgPPP1R2), Alu elements (sgAlu), and non-target control (sgNT) were based on plasmid pBA392 (a kind gift from Britt Adamson, Princeton University), a lentiviral vector for expressing sgRNA under a U6 promoter and BFP-T2A-PuromycinR under EF1*α* promoter. The target sequence (protospacer) for each sgRNA can be found in Table 2. H2B-emiRFP670 and SRRM1(2-896)-mCherry were cloned into a lentiviral vector under a UbC promoter with a GS-linker; emiRFP670 were PCR amplified from Addgene Plasmid #136571 (Matlashov et al., 2020). Plasmids were isolated from NEB Stable Competent cells using the QIAGEN Plasmid Plus Midi Kit.

**Table 1:**
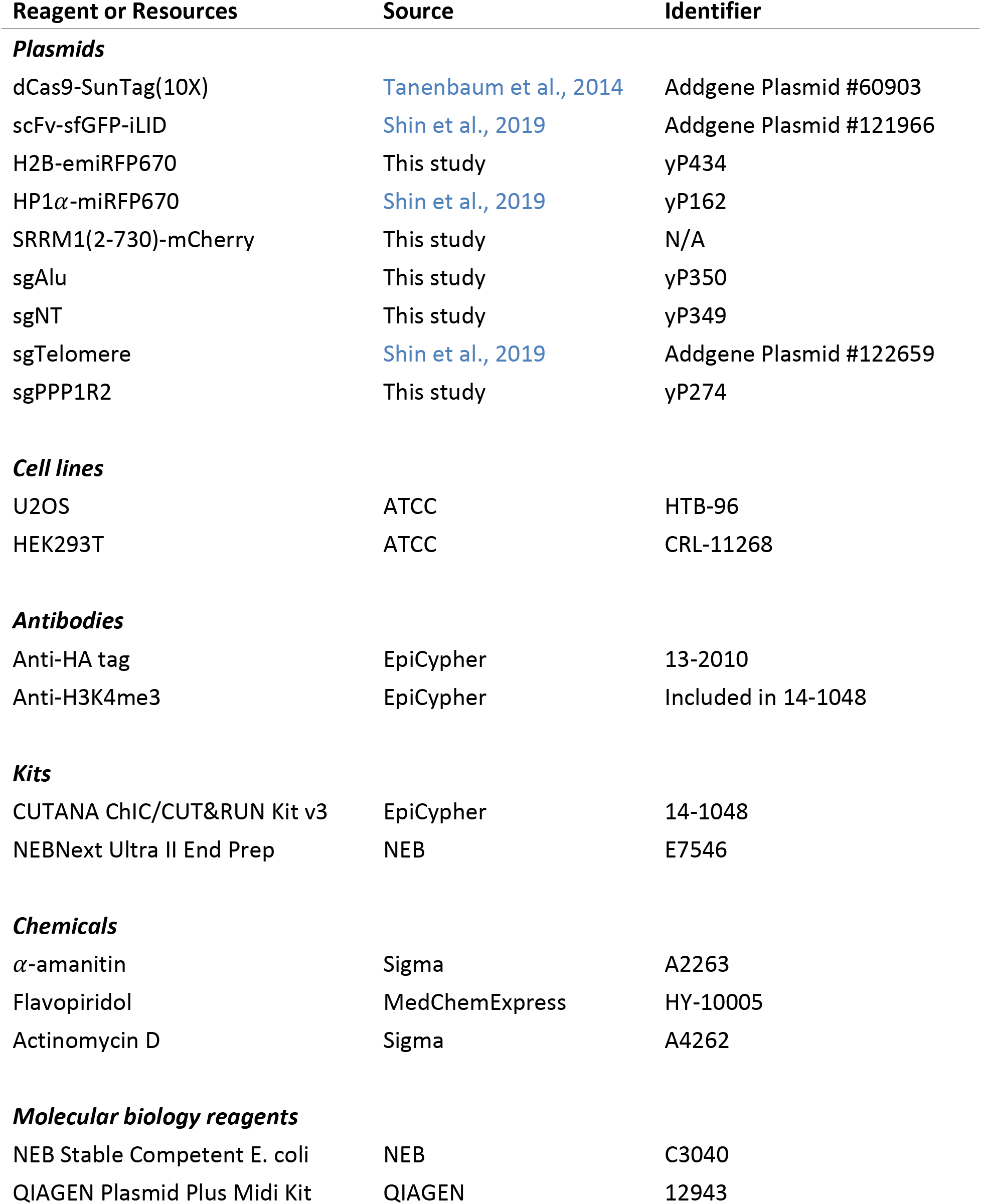
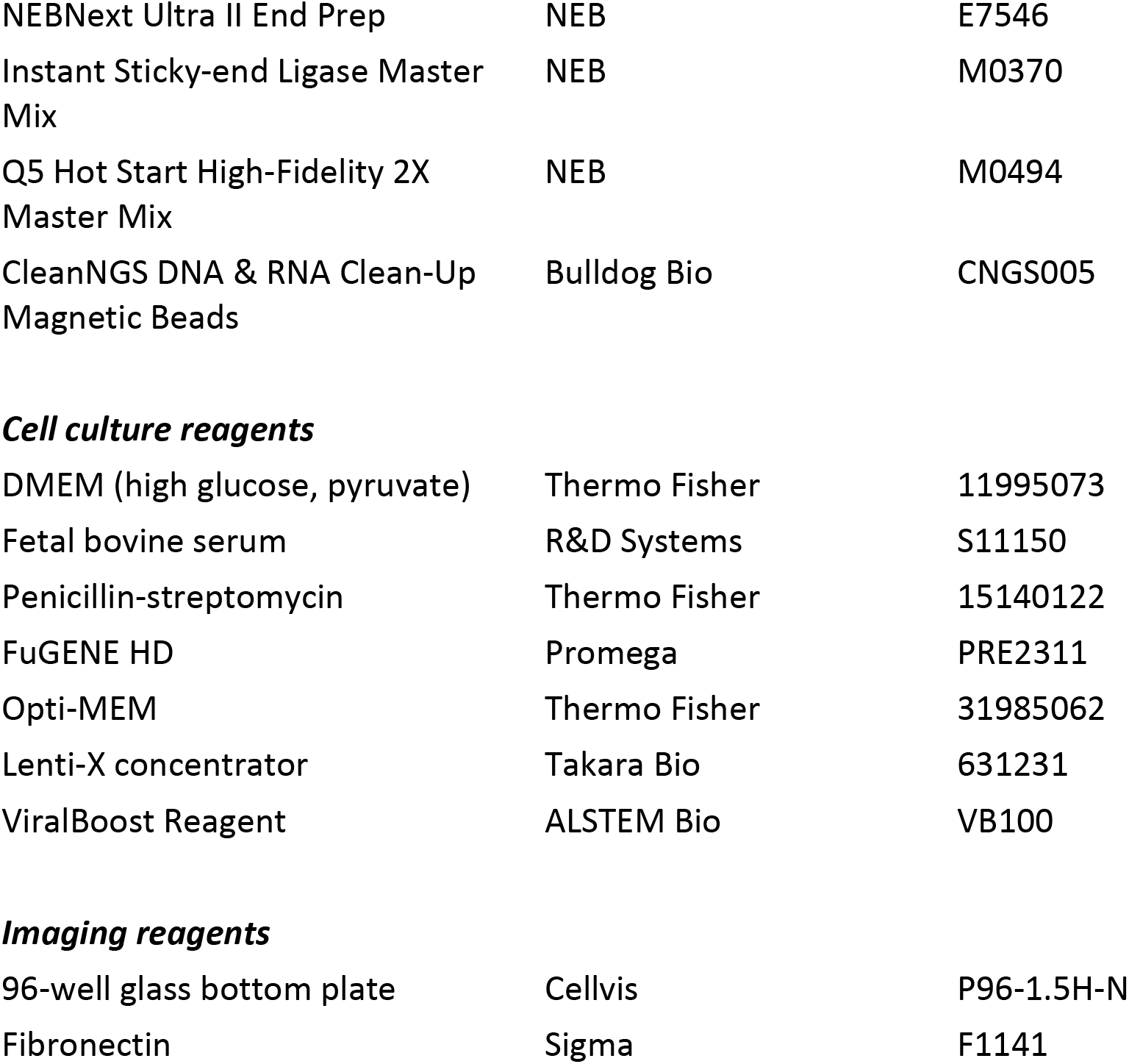
Key resources.

| Reagent or Resources | Source | Identifier |
| --- | --- | --- |
| ***Plasmids*** |  |  |
| dCas9-SunTag(10X) | Tanenbaum et al., 2014 | Addgene Plasmid #60903 |
| scFv-sfGFP-iLID | Shin et al., 2019 | Addgene Plasmid #121966 |
| H2B-emiRFP670 | This study | yP434 |
| HP1α-miRFP670 | Shin et al., 2019 | yP162 |
| SRRM1(2-730)-mCherry | This study | N/A |
| sgAlu | This study | yP350 |
| sgNT | This study | yP349 |
| sgTelomere | Shin et al., 2019 | Addgene Plasmid #122659 |
| sgPPP1R2 | This study | yP274 |
| ***Cell lines*** |  |  |
| U2OS | ATCC | HTB-96 |
| HEK293T | ATCC | CRL-11268 |
| ***Antibodies*** |  |  |
| Anti-HA tag | EpiCypher | 13-2010 |
| Anti-H3K4me3 | EpiCypher | Included in 14-1048 |
| ***Kits*** |  |  |
| CUTANA ChIC/CUT&RUN Kit v3 | EpiCypher | 14-1048 |
| NEBNext Ultra II End Prep | NEB | E7546 |
| ***Chemicals*** |  |  |
| α-amanitin | Sigma | A2263 |
| Flavopiridol | MedChemExpress | HY-10005 |
| Actinomycin D | Sigma | A4262 |
| ***Molecular biology reagents*** |  |  |
| NEB Stable Competent E. coli | NEB | C3040 |
| QIAGEN Plasmid Plus Midi Kit | QIAGEN | 12943 |
| NEBNext Ultra II End Prep | NEB | E7546 |
| Instant Sticky-end Ligase Master Mix | NEB | M0370 |
| Q5 Hot Start High-Fidelity 2X Master Mix | NEB | M0494 |
| CleanNGS DNA & RNA Clean-Up Magnetic Beads | Bulldog Bio | CNGS005 |
| ***Cell culture reagents*** |  |  |
| DMEM (high glucose, pyruvate) | Thermo Fisher | 11995073 |
| Fetal bovine serum | R&D Systems | S11150 |
| Penicillin-streptomycin | Thermo Fisher | 15140122 |
| FuGENE HD | Promega | PRE2311 |
| Opti-MEM | Thermo Fisher | 31985062 |
| Lenti-X concentrator | Takara Bio | 631231 |
| ViralBoost Reagent | ALSTEM Bio | VB100 |
| ***Imaging reagents*** |  |  |
| 96-well glass bottom plate | Cellvis | P96-1.5H-N |
| Fibronectin | Sigma | F1141 |

**Table 2:**
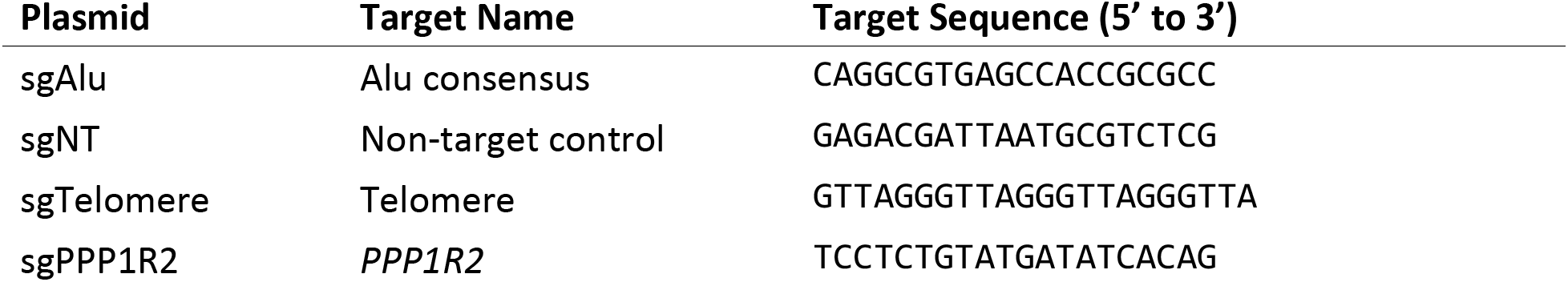
Protospacer sequences for sgRNAs used in this study.

#### Cell culture

U2OS (female) and HEK293T (female) cells were cultured in DMEM supplemented with 10% fetal bovine serum and 10 U/mL Penicillin-Streptomycin, at 37 °C and 5% CO2, in a humidified incubator.

#### Transgene expression in cultured cells

Lentiviral vectors were produced by co-transfecting HEK293T cells at ∼70% confluency in 6-well plates with transfer plasmid, pCMV-dR8.91 and pMD2.G (9:8:1 ratio) using FuGENE HD according to the manufacturer’s protocol. Each well received 3 μg of plasmid DNA and 9 μL of transfection reagent. After 12 hours, the media was replaced with fresh media containing ViralBoost reagent (at 1:500 dilution). 48, 60, and 72 hours after transfection, the media containing lentiviral particles was filtered through a 0.45 μm filter (Pall Life Sciences) and stored at 4 °C. The viral supernatant was then pooled and concentrated ten-fold to 20-fold with Lenti-X Concentrator, and then aliquots were stored at −80 °C for later use. U2OS cells were passaged and seeded at 20% confluency together with viral supernatant at the desired MOI, and fresh media was added 24 hours later. Cells were imaged at least 72 hours post-transduction.

#### Optimization for genomic imaging

U2OS cells were seeded at 20% confluency and reverse-transduced with lentiviruses expressing dCas9-HA-SunTag(10X) and scFv-sfGFP-iLID at MOI << 1. After two passages, single cells were gated to include low sfGFP signal and sorted into 96-well plates on BD FACSAria Fusion (BD Biosciences) at the Flow Cytometry Resource Facility at Princeton University. While there is a BFP marker on the dCas9 construct, the desired expression level of dCas9 is too low to allow the separation of the BFP-positive population from untransduced control cells. Single cells were then cultured and expanded for up to two weeks, passaged into 24-, 12-, and 6-well plates as necessary. Each of the successfully expanded clonal lines was split into two parts: half for maintenance, and the other half was screened for positive telomere labeling. Briefly, cells were seeded at 20% and reverse-transduced with lentivirus expressing sgRNA targeting telomere. Fresh media was added 24 hours later, and cells were passaged once before being imaged at 72-96 hours post sgRNA transduction. Positive clones identified were next screened for Alu labeling, using the half without receiving any sgRNA lentivirus and sgRNA targeting the consensus part of Alu elements. Finally, we identified clones that exhibit nucleoplasmic dCas9 signal with its CV >= 0.25 for >80% of cells to be used throughout the rest of this study. See *Quantification of dCas9 image pattern* for quantification details of this last step. The parental clones (without any sgRNA), were then expanded and frozen down for long-term storage for future experiments.

### CUT&RUN-Seq

#### Sample & library preparation and Illumina Sequencing

CUT&RUN was performed using CUTANA ChIC/CUT&RUN Kit following the manufacturer’s protocol. For CUT&RUN against dCas9 (with anti-HA tag antibody), 0.5 million U2OS cells expressing dCas9-HA-SunTag(10X) and scFv-sfGFP-iLID (described above and in Figure 1 A) each were harvested for sgAlu or sgNT conditions. For CUT&RUN against H3K4me3 (with anti-H3K4me3 antibody), 0.5 million non-transduced U2OS cells were harvested. 0.0005% digitonin was used to permeabilize the cells. All antibodies were used at 0.5 μg per reaction. 2.5 to 15 μg of purified DNA for each condition was obtained.

Illumina-compatible DNA libraries were prepared by end repair, dA-tailing, and Y-shaped adapter ligation. Briefly, NEBNext Ultra II End Prep was used to convert DNA to end-repaired DNA with 5’ phosphorylated and 3’ dA-tailed ends, and the products were cleaned up with 3X SPRI bead purification (CleanNGS beads). Next, Y-shaped adaptor-ligation was performed using Instant Sticky-end Ligase Master Mix, and adaptor-ligated products were cleaned up twice with SPRI beads (0.95X then 0.8X). Finally, DNA libraries were amplified by PCR enrichment for 12 cycles using Q5 Hot Start High-Fidelity Mastermix with dual-indexing primers incorporating the full Illumina sequencing adaptors, and cleaned up once with 0.9X SPRI. Libraries were pooled and sequenced on an Illumina Nova Seq 6000 with 150 x 150 paired-end reads.

#### CUT&RUN sequencing data processing and visualization

First, sequencing reads were trimmed using Trimmomatic (Bolger et al., 2014) v0.39 to remove sequencing adapters and low quality bases. Reads were then aligned to the human genome (GRCh38/hg38 UCSC) using STAR (Dobin et al., 2013) v2.7.8a, multi-mapping reads were filtered by removing reads with MAPQ scores < 20 using Samtools, and reads for unique alignments were finally sorted and indexed using Samtools (Danecek et al., 2021) v1.11 for visualization using pyGenomeTracks (Lopez-Delisle et al., 2021) and downstream analyses.

#### Visualizing U2OS A/B compartments

U2OS cell HiC data analyzed in this study is available in the BioStudies database (http://www.ebi.ac.uk/biostudies) under accession number: E-MTAB-8851 and source name: HiC_mOHT_rep1 (Arnould et al., 2021). To assign A and B compartments, eigenvectors were calculated using HiCExplorer (Wolff et al., 2020) with the ‘Lieberman’ method. To compare the U2OS A/B compartment annotations (aligned to hg19) to other genomic datasets generated in this study (aligned to hg38), eigenvalues were converted from hg19 to hg38 using LiftOver (UCSC) and visualized using pyGenomeTracks. Heatmaps were visualized using Juicebox (Durand et al., 2016).

#### Analyses for targeting specificity

To confirm that dCas9 localizes to Alu-containing DNA loci, we first identified which Alu repeat families match the Alu sgRNA sequence used in this study (Figure S2 B). Genomic coordinates for these sgAlu-containing Alu repeat families were then obtained from repeat masker (GRCh38/hg38 genome) (UCSC). For each subfamily, dCas9 CUT&RUN read coverage was calculated around the corresponding Alu element (+/- 500 bp) at 1-bp resolution using mapped reads with MAPQ quality scores of at least 20, using `bamCoverage` and `computeMatrix` commands from deepTools (Ramírez et al., 2016). Heatmaps and profiles were generated using `plotHeatmap`, also from deepTools.

To compare the CUT&RUN dCas9 (sgAlu and sgNT) mapped-read density with Alu-repeat annotation density genome-wide, uniquely mapped reads with quality scores of at least 20 were used. hg38 genome was binned every 1 megabase, and the read count for each bin was calculated with pair-end reads extended to match the fragment size defined by the two read mates, and normalized into counts per million (CPM) using the `bamCoverage` command from deepTools. The number of Alu annotations obtained from DFAM (Storer et al., 2021) version 3.7, for each bin was also counted using the `intersect` command from bedtools (Quinlan and Hall, 2010).

### Microscopy

#### Live cell imaging

For live cell imaging, cells were plated on the fibronectin-coated 96-well glass bottom plates and grown typically overnight before imaging.

Images shown or used for characterizing Alu-element pattern (Figure 1 B, C, Figure S1 B, and Figure 2 A, B) were acquired using a spinning disk (Yokogawa CSU-X1) confocal microscope with an EMCCD camera (Andor DU-897) and a 100X Apo TIRM objective with NA = 1.49 (MRD01991) on a Nikon Eclipse Ti body. An Okolab stage incubator was used to keep samples at 37 °C and 5% carbon dioxide during imaging sessions.

Images used for comparing H2B and Alu signals (Figure 2 C, D, and Figure S3) were acquired on a second spinning disk confocal microscope equipped with a W1 scan head (50 μm-pinhole disk), two ORCA Fusion BT back-illuminated sCMOS cameras, and a Nikon 100X Plan Apo *λ* D immersion objective (NA = 1.45), on a Nikon Ti2 body, and controlled by NIS-Elements AR software (version 5.42). A Tokai Hit stage-top incubation system was maintained at 37 °C and 5% carbon dioxide. Other images for PIV and single-particle tracking purposes were also acquired on this second microscope with protocols detailed below.

##### Live cell imaging for PIV

Simultaneous imaging of H2B- and Alu-channel by 640-nm and 488-nm lasers, respectively, yielded movies spanning 60 seconds at 2 frames per second with 200-ms exposure time for each frame, with a pixel size of 65 nm. These movies would later be analyzed using PIV (see details below). To identify nucleoli locations within each nucleus, snapshots of H2B, Alu, and BFP channels were imaged sequentially, at 640 nm, 488 nm, and 405 nm, respectively, before and after each movie, with 200-ms exposure time for each channel. Related figures: Figure 3, Figure S4, Figure S5 C, Figure 4, and Figure S6.

##### Live cell imaging for single-particle tracking

The Alu channel was imaged at enhanced spatial resolution using 2.8X SoRa magnification changer and 50 μm-pinhole SoRa SR disk for an effective pixel size of 47 nm to track individual Alu domains at a frame rate of 50 ms for 10 s continuously. Related figures: Figure S5 A, B.

#### Transcription inhibition

For transcription inhibition experiments, cells were treated with *α*-amanitin (54 μmol/L), flavopiridol (1 μmol/L), actinomycin D (1 μmol/L), or media-only (as control) for 4 to 6 hours at 37 °C prior to imaging.

#### Image analysis

All plots were generated using Python 3 with measurements obtained by software or custom scripts detailed below. Intensity-based metrics were calculated based on standardized intensity (mean-centered and standard deviation-rescaled, i.e., Z-score) to facilitate aggregation across nuclei over a range of expression levels.

#### Alu element static pattern characterization

For all static image analysis, nucleus regions were segmented based on dCas9 channel using CellPose pre-trained model for nuclei, and nucleoli regions were segmented based on BFP channel using ilastik pixel classifier. Nuclear pixels excluding nucleolar areas are referred to as nucleoplasmic regions hereon.

##### Quantification of dCas9 image pattern

The coefficient of variation (standard deviation divided by mean) of pixel intensities of the dCas9 channel in the nucleoplasmic region was calculated using measurements made by CellProfiler.

##### PCC analysis for comparing known subnuclear markers with Alu elements

For each nucleus, the Pearson correlation coefficient between dCas9 pixel intensities and subnuclear marker intensities within nucleoplasmic regions was reported by CellProfiler.

To determine the spatial correlation between Alu-rich, or A-compartment chromatin, and one of the sub-nuclear structures, we compute Pearson’s correlation coefficient (PCC) of the pixel fluorescence intensity between the two corresponding image channels (Figure 2 B). The mean of PCC values was normalized by that of the control group (sgNT).

##### Heterochromatin and euchromatin segmentation

H2B-dense region was segmented based on H2B intensity and hereon referred to as heterochromatin. The H2B image was first Gaussian-blurred (*σ* = 0.5) before binarization with a threshold equal to the 95th percentile of pixel intensity. The binary image was then subjected to morphological closing using a disk-shaped structuring element with a radius of 2 pixels. Subsequently, small areas of less than 50 pixels were removed. Euchromatic region is defined as nuclear regions excluding nucleoli or heterochromatin.

##### Alu distribution in euchromatin vs heterochromatin

The mean Z-score of Alu-channel pixel intensity within euchromatic and heterochromatic regions was reported for each nucleus.

#### Alu domain dynamics

Single particle tracking of Alu domains was performed using TrackMate (7.11.1) in Fiji with the following settings: LoG detector with threshold of 10.0, radius of 0.1, median filtered, and subpixel localization; Simple LAP tracker with max frame gap of 3, linking max distance of 0.25, gap closing max distance of 0.2, no merging allowed, and missing coordinates from closed gaps were interpolated from closest timepoints. The resulting tracking data was exported and analyzed in Python 3 using `trackpy`. Briefly, drifts were corrected by center of mass movement estimated using all tracks. Ensemble-averaged time-averaged mean square displacement (MSD) was calculated.

#### Chromatin dynamics tracking with PIV and MSND

All analyses were orchestrated using snakemake (version 7.32.4) and scripted in Python 3 and MATLAB (R2019b), and performed on Princeton University’s high-performance cluster Della.

##### Particle image velocimetry (PIV)

Calculations for PIV were done using MATLAB-based software `matpiv` (version 1.7). Briefly, 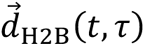, the chromatin displacement vector field for lag time τ at timepoint *t*, was calculated using the single-pass mode with a window size of 16 pixels and 75% overlap (resulting in 4-pixel spacing between vectors in either x and y directions), on H2B images taken at timepoints *t* and *t* + τ. All accessible pairs of images separated by *Δt* = τ were considered. The resulting displacement fields were filtered to keep vectors (1) whose magnitudes lie within the mean and 3 standard deviations and no more than 4 pixels, (2) where PIV correlation peak height at least 0.3, and (3) where local image signal-to-noise ratio at least 1.1. Furthermore, vectors in nucleoli, heterochromatin, or nuclear periphery regions were excluded in downstream analyses. Similarly, Alu-element displacement fields, 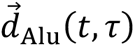, were calculated in the same way, except that Alu images and not H2B images were used.

##### Context-aware MSND analysis

To assign chromatin context (chromatin identity and environment) for each displacement vector obtained from PIV, we first Gaussian-blurring standardized context images *I*_H2B_(*t*) and *I*_Alu’_(*t*) with *σ* = 0.5, achieving approximate window-averaging with 4-pixel window size, matching the spatial resolution of displacement fields. Then, displacement fields 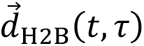 (or 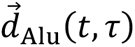) were aligned with blurred context images. Each displacement vector now would have its chromatin context values associated and be ready for further analysis.

MSND (mean square network displacement) was calculated as follows: Context values (in Z-score) were first binned (bin width = 0.25), and time-averaged ensemble-averaged mean squares of displacement vectors were calculated for each context bin. Only one context (H2B or Alu elements) was used for single-context analysis. A combination of both H2B and Alu context values was used for dual-context analysis. After aggregating data from all nuclei, context bins with less than 30 displacement vectors (or 10 for dual-context analysis) were excluded from the analysis.

## References

Arnould C, Rocher V, Finoux A-L, Clouaire T, Li K, Zhou F, Caron P, Mangeot PE, Ricci EP, Mourad R, Haber JE, Noordermeer D, Legube G. 2021. Loop extrusion as a mechanism for formation of DNA damage repair foci. Nature 590:660–665.

Bensaude O. 2011. Inhibiting eukaryotic transcription: Which compound to choose? How to evaluate its activity? Transcription 2:103–108.

Bintu L, Yong J, Antebi YE, McCue K, Kazuki Y, Uno N, Oshimura M, Elowitz MB. 2016. Dynamics of epigenetic regulation at the single-cell level. Science 351:720–724.

Bolger AM, Lohse M, Usadel B. 2014. Trimmomatic: A flexible trimmer for illumina sequence data. Bioinformatics 30:2114–2120.

Bolzer A, Kreth G, Solovei I, Koehler D, Saracoglu K, Fauth C, Müller S, Eils R, Cremer C, Speicher MR, Cremer T. 2005. Three-dimensional maps of all chromosomes in human male fibroblast nuclei and prometaphase rosettes. PLoS Biol 3:e157.

Castanon O, Smith CJ, Khoshakhlagh P, Ferreira R, Güell M, Said K, Yildiz R, Dysart M, Wang S, Thompson D, Myllykallio H, Church GM. 2020. CRISPR-mediated biocontainment. bioRxiv.

Chen B, Gilbert LA, Cimini BA, Schnitzbauer J, Zhang W, Li G-W, Park J, Blackburn EH, Weissman JS, Qi LS, Huang B. 2013. Dynamic imaging of genomic loci in living human cells by an optimized CRISPR/Cas system. Cell 155:1479–1491.

Chen Y, Zhang Y, Wang Y, Zhang L, Brinkman EK, Adam SA, Goldman R, Steensel B van, Ma J, Belmont AS. 2018. Mapping 3D genome organization relative to nuclear compartments using TSA-Seq as a cytological ruler. J Cell Biol 217:4025–4048.

Danecek P, Bonfield JK, Liddle J, Marshall J, Ohan V, Pollard MO, Whitwham A, Keane T, McCarthy SA, Davies RM, Li H. 2021. Twelve years of SAMtools and BCFtools. Gigascience 10.

Deininger P. 2011. Alu elements: Know the SINEs. Genome Biol 12:236.

Dixon JR, Jung I, Selvaraj S, Shen Y, Antosiewicz-Bourget JE, Lee AY, Ye Z, Kim A, Rajagopal N, Xie W, Diao Y, Liang J, Zhao H, Lobanenkov VV, Ecker JR, Thomson JA, Ren B. 2015. Chromatin architecture reorganization during stem cell differentiation. Nature 518:331–336.

Dobin A, Davis CA, Schlesinger F, Drenkow J, Zaleski C, Jha S, Batut P, Chaisson M, Gingeras TR. 2013. STAR: Ultrafast universal RNA-seq aligner. Bioinformatics 29:15–21.

Durand NC, Robinson JT, Shamim MS, Machol I, Mesirov JP, Lander ES, Aiden EL. 2016. Juicebox provides a visualization system for Hi-C contact maps with unlimited zoom. Cell Syst 3:99–101.

Eeftens JM, Kapoor M, Michieletto D, Brangwynne CP. 2021. Polycomb condensates can promote epigenetic marks but are not required for sustained chromatin compaction. Nat Commun 12:5888.

Gu B, Swigut T, Spencley A, Bauer MR, Chung M, Meyer T, Wysocka J. 2018. Transcription-coupled changes in nuclear mobility of mammalian cis-regulatory elements. Science 359:1050– 1055.

Guntas G, Hallett RA, Zimmerman SP, Williams T, Yumerefendi H, Bear JE, Kuhlman B. 2015. Engineering an improved light-induced dimer (iLID) for controlling the localization and activity of signaling proteins. Proc Natl Acad Sci U S A 112:112–117.

Kimura H, Sugaya K, Cook PR. 2002. The transcription cycle of RNA polymerase II in living cells. J Cell Biol 159:777–782.

Kölbl AC, Weigl D, Mulaw M, Thormeyer T, Bohlander SK, Cremer T, Dietzel S. 2012. The radial nuclear positioning of genes correlates with features of megabase-sized chromatin domains. Chromosome Res 20:735–752.

Ku H, Park G, Goo J, Lee J, Park TL, Shim H, Kim JH, Cho W-K, Jeong C. 2022. Effects of transcription-dependent physical perturbations on the chromosome dynamics in living cells. Front Cell Dev Biol 10:822026.

Lander ES, Linton LM, Birren B, Nusbaum C, Zody MC, Baldwin J, Devon K, Dewar K, Doyle M, FitzHugh W, Funke R, Gage D, Harris K, Heaford A, Howland J, Kann L, Lehoczky J, LeVine R, McEwan P, McKernan K, Meldrim J, Mesirov JP, Miranda C, Morris W, Naylor J, Raymond C, Rosetti M, Santos R, Sheridan A, Sougnez C, Stange-Thomann Y, Stojanovic N, Subramanian A, Wyman D, Rogers J, Sulston J, Ainscough R, Beck S, Bentley D, Burton J, Clee C, Carter N, Coulson A, Deadman R, Deloukas P, Dunham A, Dunham I, Durbin R, French L, Grafham D, Gregory S, Hubbard T, Humphray S, Hunt A, Jones M, Lloyd C, McMurray A, Matthews L, Mercer S, Milne S, Mullikin JC, Mungall A, Plumb R, Ross M, Shownkeen R, Sims S, Waterston RH, Wilson RK, Hillier LW, McPherson JD, Marra MA, Mardis ER, Fulton LA, Chinwalla AT, Pepin KH, Gish WR, Chissoe SL, Wendl MC, Delehaunty KD, Miner TL, Delehaunty A, Kramer JB, Cook LL, Fulton RS, Johnson DL, Minx PJ, Clifton SW, Hawkins T, Branscomb E, Predki P, Richardson P, Wenning S, Slezak T, Doggett N, Cheng JF, Olsen A, Lucas S, Elkin C, Uberbacher E, Frazier M, Gibbs RA, Muzny DM, Scherer SE, Bouck JB, Sodergren EJ, Worley KC, Rives CM, Gorrell JH, Metzker ML, Naylor SL, Kucherlapati RS, Nelson DL, Weinstock GM, Sakaki Y, Fujiyama A, Hattori M, Yada T, Toyoda A, Itoh T, Kawagoe C, Watanabe H, Totoki Y, Taylor T, Weissenbach J, Heilig R, Saurin W, Artiguenave F, Brottier P, Bruls T, Pelletier E, Robert C, Wincker P, Smith DR, Doucette-Stamm L, Rubenfield M, Weinstock K, Lee HM, Dubois J, Rosenthal A, Platzer M, Nyakatura G, Taudien S, Rump A, Yang H, Yu J, Wang J, Huang G, Gu J, Hood L, Rowen L, Madan A, Qin S, Davis RW, Federspiel NA, Abola AP, Proctor MJ, Myers RM, Schmutz J, Dickson M, Grimwood J, Cox DR, Olson MV, Kaul R, Raymond C, Shimizu N, Kawasaki K, Minoshima S, Evans GA, Athanasiou M, Schultz R, Roe BA, Chen F, Pan H, Ramser J, Lehrach H, Reinhardt R, McCombie WR, Bastide M de la, Dedhia N, Blöcker H, Hornischer K, Nordsiek G, Agarwala R, Aravind L, Bailey JA, Bateman A, Batzoglou S, Birney E, Bork P, Brown DG, Burge CB, Cerutti L, Chen HC, Church D, Clamp M, Copley RR, Doerks T, Eddy SR, Eichler EE, Furey TS, Galagan J, Gilbert JG, Harmon C, Hayashizaki Y, Haussler D, Hermjakob H, Hokamp K, Jang W, Johnson LS, Jones TA, Kasif S, Kaspryzk A, Kennedy S, Kent WJ, Kitts P, Koonin EV, Korf I, Kulp D, Lancet D, Lowe TM, McLysaght A, Mikkelsen T, Moran JV, Mulder N, Pollara VJ, Ponting CP, Schuler G, Schultz J, Slater G, Smit AF, Stupka E, Szustakowki J, Thierry-Mieg D, Thierry-Mieg J, Wagner L, Wallis J, Wheeler R, Williams A, Wolf YI, Wolfe KH, Yang SP, Yeh RF, Collins F, Guyer MS, Peterson J, Felsenfeld A, Wetterstrand KA, Patrinos A, Morgan MJ, Jong P de, Catanese JJ, Osoegawa K, Shizuya H, Choi S, Chen YJ, Szustakowki J, International Human Genome Sequencing Consortium. 2001. Initial sequencing and analysis of the human genome. Nature 409:860–921.

Lieberman-Aiden E, Berkum NL van, Williams L, Imakaev M, Ragoczy T, Telling A, Amit I, Lajoie BR, Sabo PJ, Dorschner MO, Sandstrom R, Bernstein B, Bender MA, Groudine M, Gnirke A, Stamatoyannopoulos J, Mirny LA, Lander ES, Dekker J. 2009. Comprehensive mapping of long-range interactions reveals folding principles of the human genome. Science 326:289–293.

Liu WM, Maraia RJ, Rubin CM, Schmid CW. 1994. Alu transcripts: Cytoplasmic localisation and regulation by DNA methylation. Nucleic Acids Res 22:1087–1095.

Lopez-Delisle L, Rabbani L, Wolff J, Bhardwaj V, Backofen R, Grüning B, Ramırez F, Manke T. 2021. pyGenomeTracks: Reproducible plots for multivariate genomic datasets. Bioinformatics 37:422–423.

Lu JY, Chang L, Li T, Wang T, Yin Y, Zhan G, Han X, Zhang K, Tao Y, Percharde M, Wang L, Peng Q, Yan P, Zhang H, Bi X, Shao W, Hong Y, Wu Z, Ma R, Wang P, Li W, Zhang J, Chang Z, Hou Y, Zhu B, Ramalho-Santos M, Li P, Xie W, Na J, Sun Y, Shen X. 2021. Homotypic clustering of L1 and B1/Alu repeats compartmentalizes the 3D genome. Cell Res 31:613–630.

Lu JY, Shao W, Chang L, Yin Y, Li T, Zhang H, Hong Y, Percharde M, Guo L, Wu Z, Liu L, Liu W, Yan P, Ramalho-Santos M, Sun Y, Shen X. 2020. Genomic repeats categorize genes with distinct functions for orchestrated regulation. Cell Rep 30:3296–3311.e5.

Maeshima K, Iida S, Shimazoe MA, Tamura S, Ide S. 2023. Is euchromatin really open in the cell? Trends Cell Biol.

Marchal C, Sima J, Gilbert DM. 2019. Control of DNA replication timing in the 3D genome. Nat Rev Mol Cell Biol 20:721–737.

Matlashov ME, Shcherbakova DM, Alvelid J, Baloban M, Pennacchietti F, Shemetov AA, Testa I, Verkhusha VV. 2020. A set of monomeric near-infrared fluorescent proteins for multicolor imaging across scales. Nat Commun 11:239.

Miron E, Oldenkamp R, Brown JM, Pinto DMS, Xu CS, Faria AR, Shaban HA, Rhodes JDP, Innocent C, Ornellas S de, Hess HF, Buckle V, Schermelleh L. 2020. Chromatin arranges in chains of mesoscale domains with nanoscale functional topography independent of cohesin. Sci Adv 6.

Miura H, Hiratani I. 2022. Cell cycle dynamics and developmental dynamics of the 3D genome: Toward linking the two timescales. Curr Opin Genet Dev 73:101898.

Miura M, Chen H. 2020. CUT&RUN detects distinct DNA footprints of RNA polymerase II near the transcription start sites. Chromosome Res 28:381–393.

Nagashima R, Hibino K, Ashwin SS, Babokhov M, Fujishiro S, Imai R, Nozaki T, Tamura S, Tani T, Kimura H, Shribak M, Kanemaki MT, Sasai M, Maeshima K. 2019. Single nucleosome imaging reveals loose genome chromatin networks via active RNA polymerase II. J Cell Biol 218:1511– 1530.

Nozaki T, Shinkai S, Ide S, Higashi K, Tamura S, Shimazoe MA, Nakagawa M, Suzuki Y, Okada Y, Sasai M, Onami S, Kurokawa K, Iida S, Maeshima K. 2023. Condensed but liquid-like domain organization of active chromatin regions in living human cells. Science Advances 9:eadf1488.

Ochiai H, Sugawara T, Yamamoto T. 2015. Simultaneous live imaging of the transcription and nuclear position of specific genes. Nucleic Acids Res 43:e127.

Paulson KE, Schmid CW. 1986. Transcriptional inactivity of alu repeats in HeLa cells. Nucleic Acids Res 14:6145–6158.

Quinlan AR, Hall IM. 2010. BEDTools: A flexible suite of utilities for comparing genomic features. Bioinformatics 26:841–842.

Ramírez F, Ryan DP, Grüning B, Bhardwaj V, Kilpert F, Richter AS, Heyne S, Dündar F, Manke T. 2016. deepTools2: A next generation web server for deep-sequencing data analysis. Nucleic Acids Res 44:W160–5.

Rowley MJ, Nichols MH, Lyu X, Ando-Kuri M, Rivera ISM, Hermetz K, Wang P, Ruan Y, Corces VG. 2017. Evolutionarily conserved principles predict 3D chromatin organization. Mol Cell 67:837–852.e7.

Sato Y, Mukai M, Ueda J, Muraki M, Stasevich TJ, Horikoshi N, Kujirai T, Kita H, Kimura T, Hira S, Okada Y, Hayashi-Takanaka Y, Obuse C, Kurumizaka H, Kawahara A, Yamagata K, Nozaki N, Kimura H. 2013. Genetically encoded system to track histone modification in vivo. Sci Rep 3:2436.

Saxton MN, Morisaki T, Krapf D, Kimura H, Stasevich TJ. 2023. Live-cell imaging uncovers the relationship between histone acetylation, transcription initiation, and nucleosome mobility. Sci Adv 9:eadh4819.

Shaban HA, Barth R, Recoules L, Bystricky K. 2020. Hi-D: Nanoscale mapping of nuclear dynamics in single living cells. Genome Biol 21:95.

Shin S, Cho HW, Shi G, Thirumalai D. 2023. Transcription-induced active forces suppress chromatin motion by inducing a transient disorder-to-order transition. arXiv 122:19a.

Shin Y, Chang Y-C, Lee DSW, Berry J, Sanders DW, Ronceray P, Wingreen NS, Haataja M, Brangwynne CP. 2019. Liquid nuclear condensates mechanically sense and restructure the genome. Cell 176:1518.

Skene PJ, Henikoff S. 2017. An efficient targeted nuclease strategy for high-resolution mapping of DNA binding sites. Elife 6.

Solovei I, Kreysing M, Lanctôt C, Kösem S, Peichl L, Cremer T, Guck J, Joffe B. 2009. Nuclear architecture of rod photoreceptor cells adapts to vision in mammalian evolution. Cell 137:356– 368.

Storer J, Hubley R, Rosen J, Wheeler TJ, Smit AF. 2021. The dfam community resource of transposable element families, sequence models, and genome annotations. Mob DNA 12:2.

Su J-H, Zheng P, Kinrot SS, Bintu B, Zhuang X. 2020. Genome-Scale imaging of the 3D organization and transcriptional activity of chromatin. Cell 182:1641–1659.e26.

Talbert PB, Meers MP, Henikoff S. 2019. Old cogs, new tricks: The evolution of gene expression in a chromatin context. Nat Rev Genet 20:283–297.

Tanenbaum ME, Gilbert LA, Qi LS, Weissman JS, Vale RD. 2014. A protein-tagging system for signal amplification in gene expression and fluorescence imaging. Cell 159:635–646.

Wolff J, Rabbani L, Gilsbach R, Richard G, Manke T, Backofen R, Grüning BA. 2020. Galaxy HiCExplorer 3: A web server for reproducible Hi-C, capture Hi-C and single-cell Hi-C data analysis, quality control and visualization. Nucleic Acids Res 48:W177–W184.

Xie L, Dong P, Chen X, Hsieh T-HS, Banala S, De Marzio M, English BP, Qi Y, Jung SK, Kieffer-Kwon K-R, Legant WR, Hansen AS, Schulmann A, Casellas R, Zhang B, Betzig E, Lavis LD, Chang HY, Tjian R, Liu Z. 2020. 3D ATAC-PALM: Super-resolution imaging of the accessible genome. Nat Methods 17:430–436.

Xie L, Liu Z. 2021. Single-cell imaging of genome organization and dynamics. Mol Syst Biol 17:e9653.

Zheng H, Xie W. 2019. The role of 3D genome organization in development and cell differentiation. Nat Rev Mol Cell Biol 20:535–550.

Zidovska A, Weitz DA, Mitchison TJ. 2013. Micron-scale coherence in interphase chromatin dynamics. Proc Natl Acad Sci U S A 110:15555–15560.

